# Fluctuating salinity during development impacts fish productivity

**DOI:** 10.1101/2024.02.01.578515

**Authors:** Meng-Han Joseph Chung, Daniel W. A. Noble, Rebecca J. Fox, Lauren M. Harrison, Michael D. Jennions

## Abstract

Climate change is elevating salinity levels in many freshwater systems, and more erratic rainfall is increasing variation in salinity. Consequently, many species now experience more extreme developmental environments. Resultant shifts in developmental trajectories could change key life history traits that persist into adulthood. To date, however, how variation in salinity affects the life histories of freshwater species has been neglected despite its implications for fisheries. We ran a large-scale experiment with a global pest, the mosquitofish (*Gambusia holbrooki*), and manipulated the salinity experienced by juveniles: freshwater (0‰), stable salinity (10‰) or fluctuating salinity (0-20‰; mean = 10 ‰). Fish developing in stable, high salinity grew faster and matured earlier, albeit with a decline in male telomeres and female gut development. Stable high salinity resulted in larger adult body size in females, but not males, which increased female fecundity. Conversely, fluctuations in salinity induced fish to grow more slowly and lowered female fecundity. Crucially, several of the long-term effects of salinity fluctuations were sex-specific, more adversely affecting females than males. We highlight that environmental variability alters an organism’s vulnerability to stressors, with implications that should be considered if we wish to understand the impact of climate change on population dynamics.

## INTRODUCTION

Climate change is a major driver of species extinction in the 21^st^ century^1, 2^, partially because populations are experiencing more variable environmental conditions^3,4^. Rapidly fluctuating environments create shorter periods of more intense stress than is typical in stable environments, implying that variability is harmful^5^. For aquatic organisms, climate change is not only elevating water temperature but, perhaps more importantly, it is changing salinity.

Fluctuations in salinity occur due to variation in precipitation, tidal flow, the occurrence of flooding, and seawater intrusion into freshwater bodies^6-8^. On top of climate-driven changes, many human activities amplify the magnitude and extent of natural fluctuations in salinity by altering water run-off (e.g., land clearing) and adding salt to freshwater (e.g., industrial effluents)^7-9^. For example, wastewater from shale gas operations has made salinity levels seven times higher than that of seawater in the U.S.A.^10^, with a similar issue reported in the Amazon due to oil extraction^11^. Currently, salinization is a pressing global environmental problem^12-14^, affecting over 30% of freshwater ecosystems^8^. It is a serious threat to freshwater biodiversity^13^. For example, higher salinity in the Aral Sea led to its freshwater ecosystem being replaced by a marine ecosystem, collapsing local fisheries^15^.

Environmental variability is predicted to affect how animals allocate limited resources to key processes, creating differences in the ‘pace of life’^16^. Under stressful conditions, individuals sometimes adopt a ‘fast’ pace of life, characterized by rapid growth, earlier maturation, and an earlier onset and faster rate of reproduction. This often comes at the cost of accelerated senescence and reduced longevity^17, 18^. This ‘fast’ strategy allows individuals to produce more offspring in a shorter period, which is consistent with a life history that has adapted to high mortality such that the future is ‘discounted’^19^. Conversely, greater variability also involves periods of reduced stress that might promote recovery^20^. Exposure to a stressful environment might therefore favor a ‘slow’ pace of life if greater investment in self-maintenance can sufficiently improve survival, even if this diverts resources away from growth and lowers the rate of reproduction^21^. Ultimately, the net effect of greater environmental variability on fitness is uncertain, and life history theory struggles to make clear predictions^22^.

Natural selection favors life-history strategies that optimize allocation to growth, reproduction and self-maintenance, enabling organisms to persist in diverse habitats^23^. In stable environments, selection is expected to promote local adaptation of the life history^24^. Optimal life history strategies partly depend on how environmental factors affect extrinsic rates of mortality at different life stages^25^ (e.g., juveniles versus adults), and how shifts in resource allocation moderate the risk of mortality^26^. The effect of environmental variability on life-histories is also likely to differ between the sexes. Males and females usually experience sex-specific selection on life-history strategies that determine their fitness^22, 27^. Females often benefit from ongoing investment into growth because fecundity increases rapidly with body size^28^. In contrast, males in many polyandrous species invest heavily into sexually selected traits (e.g., weapons, courtship) to improve access to mates, with minimal growth after maturation^29^, faster senescence^30^ and greater mortality^31^. To date, we know little about the extent of sex differences in plastic (or evolved) adjustment of life histories in response to the level of stressors. More importantly, we know almost nothing about how greater short-term variability in environmental stressors affects life history shifts in each sex despite an understanding being key to predicting population viability in a changing climate.

To date, no study has examined how fluctuating versus constant levels of high salinity during development affect fish life histories. This is an oversight as the richness of freshwater species declines dramatically when salinity levels exceed ±10‰^32, 33^, which is now occurring in many places due to anthropogenic factors^5^. To test the effect on life history traits of fluctuating versus constant salinity during development, we ran a large-scale experiment on the freshwater mosquitofish (*Gambusia holbrooki*). This is one of the most invasive fish worldwide^34^, known for its aggressive behavior towards native fish^35, 36^. Mosquitofish are hard to eradicate, partly due to their ability to tolerate environmental stressors^34^. However, established populations are susceptible to changes in salinity^37, 38^ due to the drying of streams, proximity to estuaries, and salinization after land clearing^39-41^. Here, we investigate how both the level and variability in salinity during development affects life history traits. *G. holbrooki* are a model species to more broadly test how freshwater fish might respond to increased and more variable salinity.

## RESULTS

We randomly assigned newborn mosquitofish into three treatments: freshwater (control), stable salinity (10‰), or fluctuating salinity environments (0-20‰, mean = 10‰). Fish in the fluctuating treatment experienced salinity changes of 10‰ every two days. All fish were reared individually and transferred to freshwater upon maturation to test how the developmental environment affects adult life history traits in a common garden setting. We measured a suite of adult traits (growth, reproduction, survival, and aspects of self-maintenance), both when fish were young (shortly after maturity) and old (after another 12- weeks during which fish mated and bred). We collected data on both females (*n* = 606) and males (*n* = 567) to test for sex differences in how fluctuations in salinity affect life histories.

### Salinity and its fluctuations had sex-specific effects on growth

The developmental environment significantly affected juvenile growth (Supplementary Fig. 1). Growth was fastest with stable salinity and slowest with fluctuating salinity (Supplementary Table 1*A*). Consequently, juvenile size differed significantly between all three environments from the third week after birth onward (pairwise: all *P* < 0.001).

Females matured significantly later when they developed in a fluctuating rather than stable salinity environment (Fig. 1A), with females from freshwater taking an intermediate time to mature (pairwise: all *P* < 0.001). There was, however, no significant difference in female size at maturity between the two salinity environments (*P* = 0.202; Supplementary Table 3).

**Fig 1.**
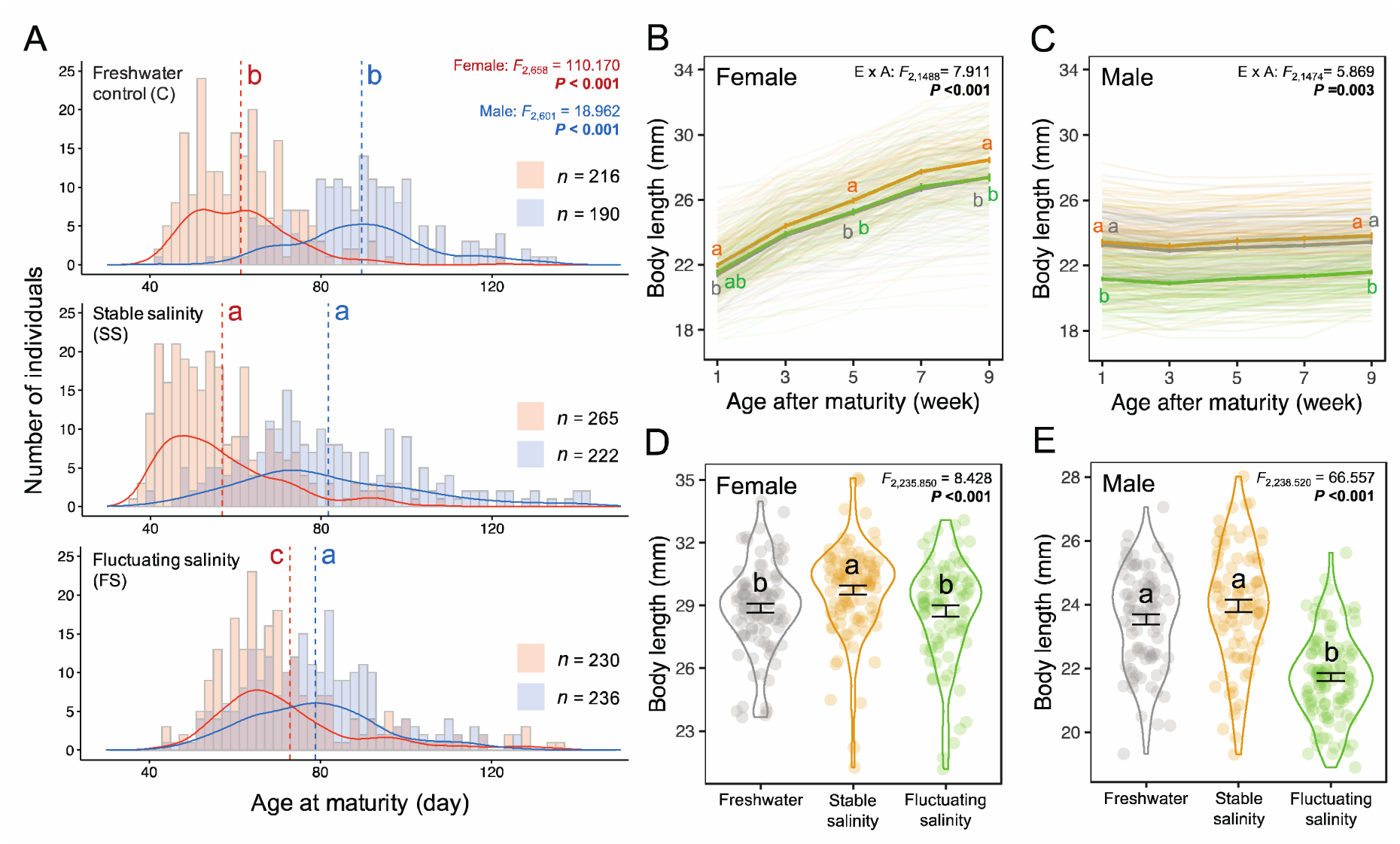
Effects of developmental environment on (A) age at maturity, (B, C) adult growth and (D, E) body size at the end of the experiment for both sexes. Days to mature (A) of females (red) and males (blue) are shown individually for each environment in a combination of histogram and density plots (mean = dashed line). Line plots (B, C) show individual growth (thin lines) and mean growth (thick lines) in each environment in the first eight weeks after returning to freshwater. Violin plots (D, E) show the body length at the end of the mating period (Week 13 following maturation for females; Week 15 for males). Colors represent three environments: freshwater control (grey); stable salinity (orange); fluctuating salinity (green). E×A indicates an environment-by-age interaction. Different letters represent significant differences among environments from Tukey’s tests. For adult growth, letters are only shown at the start and end of the adult period and when the significance levels changed; no letter at an age indicates that the significant difference seen at the previous age persists (see Supplementary Table 1). Error bars represent mean ± *SE*.

Females from the stable (*P* = 0.007), but not the fluctuating (*P* = 0.326), salinity environment were significantly larger than those from the freshwater environment. Females continued to grow after maturation: five weeks later those from the stable salinity environment were the largest (Fig. 1B, D).

Unlike females, males that developed in freshwater matured significantly later than those from either saline environment (pairwise: both *P* < 0.001) (Fig. 1A), but there was no effect of fluctuations in salinity (*P* = 0.168). Males from the fluctuating salinity environment were significantly smaller at maturity than those from the stable salinity or freshwater environments (both *P* < 0.001; Supplementary Table 3), with no size difference between the latter two (*P* = 0.168). Adult males barely grew, so males from the fluctuating salinity environment remained significantly smaller throughout adulthood (all *P* < 0.001; Fig. 1C, E).

### Salinity and its fluctuations had no effect on survival, but salinity had sex-specific effects on senescence and somatic traits

The developmental environment had no effect on female or male survival (Supplementary Fig. 2), but this does not preclude other costs of higher and/or fluctuating salinity. Developmental environments can affect self-maintenance traits (e.g., immune responses^42^) that moderate how energy is allocated to reproduction. For example, fluctuating salinity is expected to pose osmoregulatory challenges^43, 44^, which could lower investment in other fitness-enhancing traits.

We measured relative telomere length (RTL) in a set of young and old adults. Telomeres are repetitive sequences of noncoding DNA at chromosome ends, protecting coding sequences from attrition^45^. Telomere shortening is often linked to ageing^46^, making RTL a potential indicator of the rate of senescence and lifespan^47, 48^. The developmental environment affected female RTL, but this was moderated by female age (i.e., an interaction: LMM, *F* = 3.998, *P* = 0.021). There was, however, no significant environmental effect on RTL when looking separately at young (LMM, *F* = 1.602, *P* = 0.210) and old females (LMM, *F* = 2.429, *P* = 0.097). The interaction was driven by females from freshwater having the shortest RTL when young, but the longest RTL when old (Supplementary Fig. 3A), suggesting a slower rate of telomere shortening for females that developed in freshwater. Males showed a different pattern: the developmental environment affected RTL (LMM, *F* = 7.908, *P* < 0.001), irrespective of age (*F* = 1.451, *P* = 0.238). Males from the stable salinity environment had a significantly shorter RTL than those from freshwater (pairwise: *P* < 0.001) (Supplementary Fig. 4A), while the RTL of males from the fluctuating salinity environment was intermediate (both *P* > 0.064). Age (days since birth) and body size had no effect on female or male RTL (Supplementary Tables 4-7).

The gut plays an important role in salinity acclimation through ion uptake that reduces gut osmolarity to facilitate water uptake^49, 50^. A longer gut potentially enhances absorption^51^, but intestinal cells have a high turnover rate^52^, suggesting that a longer gut imposes maintenance costs^53, 54^. For females, the developmental environment affected their gut length after controlling for body size (LMM, *F* = 11.874, *P* < 0.001). Females from the stable salinity environment had a significantly shorter gut than those from either the freshwater or fluctuating salinity environments (pairwise: both *P* < 0.001), with no difference between the latter two environments (*P* = 0.855) (Supplementary Fig. 3B). Males showed a different pattern: there was no effect of the developmental environment on relative gut length (LMM, *F* = 0.482, *P* = 0.618).

Finally, we measured immune responses using a phytohemagglutinin injection assay^55^. There were no sex-specific effects. Neither the developmental environment, adult age, their interaction, nor body size (Supplementary Tables 4-7) affected immunocompetence.

### Females that developed under fluctuating salinity had a lower reproductive output

Mosquitofish are livebearers that gives birth to well-developed offspring. We dissected a subset of newly mature females to analyze the size and number of unfertilized eggs. The remaining females were paired with randomly selected stock males in a common garden freshwater environment for 12 weeks to assess overall offspring production. After this period, we dissected older females to quantify the number of eggs and embryos. This enabled us to test for developmental environmental effects on female fecundity while controlling for any effects of the developmental environment on male reproduction.

The developmental environment experienced and age interacted to determine total egg number (GLMM, χ^2^ = 6.642, *P* = 0.036). Young females produced a similar number of eggs irrespective of the environment (GLMM, χ^2^ = 0.193, *P* = 0.908). However, old females from the fluctuating salinity environment produced significantly fewer eggs than those from the stable salinity environment (pairwise: *P* < 0.001), while neither differed from females from freshwater (both *P* > 0.078) (Fig. 2*A*). Egg size was affected by the developmental environment (LMM, *F* = 12.642, *P* < 0.001) (Fig. 2*B*). Females from the fluctuating salinity environment had significantly smaller eggs than those of females from the stable salinity or freshwater environments (pairwise: both *P* < 0.002), with no difference between the latter two (*P* = 0.855).

**Fig 2.**
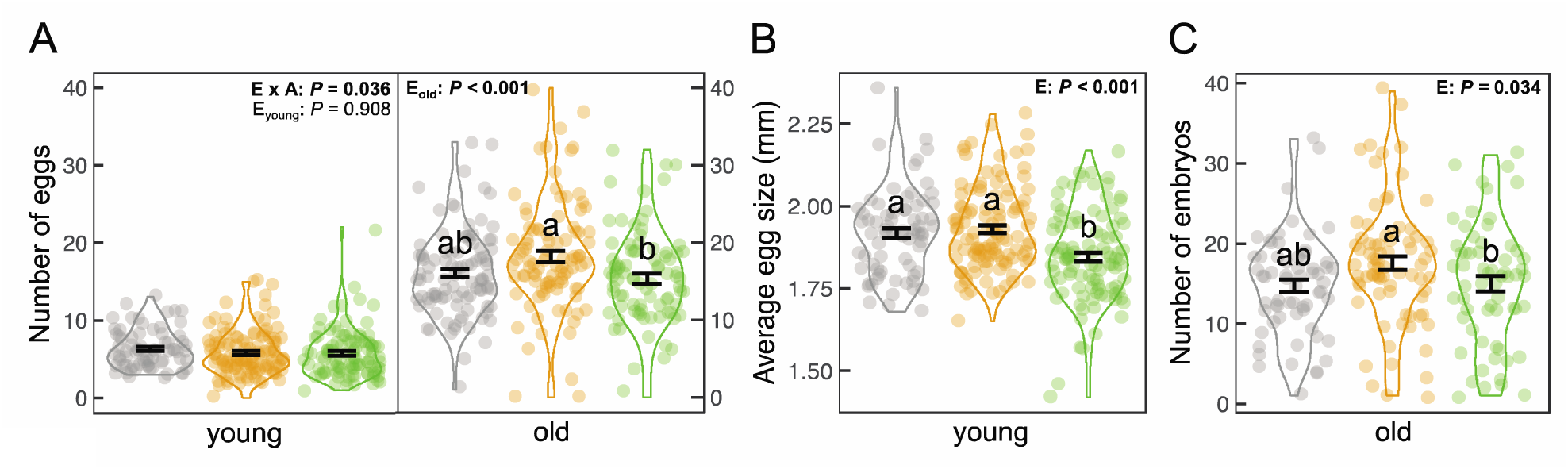
Effect of development environment and adult age on female fecundity. Colors indicate the environments: freshwater (F; grey); stable salinity (SS; orange); fluctuating salinity (FS; green). (A) total egg number (*young* F: *n* = 72; SS: *n* = 118, FS: *n* = 100; *old* F: *n* = 99; SS: *n* = 98; FS: *n* = 87), (B) egg size of young females (F: *n* = 72; SS: *n* = 114; FS: *n* = 100), (C) embryo number of old females (F: *n* = 64; SS: *n* = 78; FS: *n* = 62) are presented with the significance levels of environment (E), age (A) and their interaction (E×A) using bold font. Given a significant interaction, the environment effect was shown separately for each age class. Figures show mean ± *SE* and only indicate the conditional part of the results for (C) embryo number. Different letters represent significant differences between environments based on Tukey’s tests (see Supplementary Table 4 for details).

The developmental environment affected the likelihood of carrying embryos (GLMM, χ^2^ = 6.145, *P* = 0.046) and, if present, the number of embryos (χ^2^ = 6.755, *P* = 0.034). The likelihood of carrying embryos did not differ between the two saline environments (pairwise: *P* = 0.290), but females from the stable salinity environment carried more embryos than females from the fluctuating salinity environment (*P* = 0.042) (Fig. 2*C*). On the other hand, females from the stable salinity environment were more likely to have embryos than those from the freshwater environment (*P* = 0.037), but the number of embryos did not differ between the freshwater and stable salinity environments (*P* = 0.161).

The likelihood of giving birth did not depend on the developmental environment (GLMM, χ^2^ = 0.759, *P* = 0.684), but the total number of offspring (i.e., births and embryos) did (GLMM, χ = 7.865, *P* = 0.020). Females from the fluctuating salinity environment had marginally non-significantly fewer offspring than those from the stable salinity (pairwise: *P* = 0.051). Females from the stable salinity environment produced significantly more offspring than those from the freshwater environment (*P* = 0.047).

Larger females were more fecund (Supplementary Fig. 6). After controlling for female size, the developmental environment no longer influenced the number of eggs or embryos, or total number of offspring (Supplementary Table 6). The effects of developmental environment on fecundity therefore seem to be mediated by its effect on female size.

### Salinity enhanced the mating effort of older males, but salinity fluctuations lowered sperm count

Differences in population growth can also be influenced by male reproductive effort (e.g., sexual harassment lowers female fecundity^56^). We paired focal males with random stock females in freshwater for 12 weeks to test the effect of the developmental environment on male mating behavior (time spent pursuing females, number of mating attempts, likelihood of success) and ejaculate traits (sperm count, velocity). These traits influence male reproduction success^57, 58^.

The developmental environment and male age interacted to influence the number of mating attempts (GLMM, χ^2^ = 8.498, *P* = 0.014) (Fig. 3*A*). Old males from both salinity environments (pairwise: *P* = 0.994) made significantly more mating attempts than those from freshwater (both *P* < 0.015) (GLMM, χ^2^ = 11.367, *P* = 0.003). In contrast, there was no environmental effect for young males (GLMM, χ^2^ = 0.986, *P* = 0.611). The developmental environment had no effect on the likelihood of a successful mating attempt (Supplementary Table 5) or the time spent pursuing females (Fig. 3*B*).

**Fig 3.**
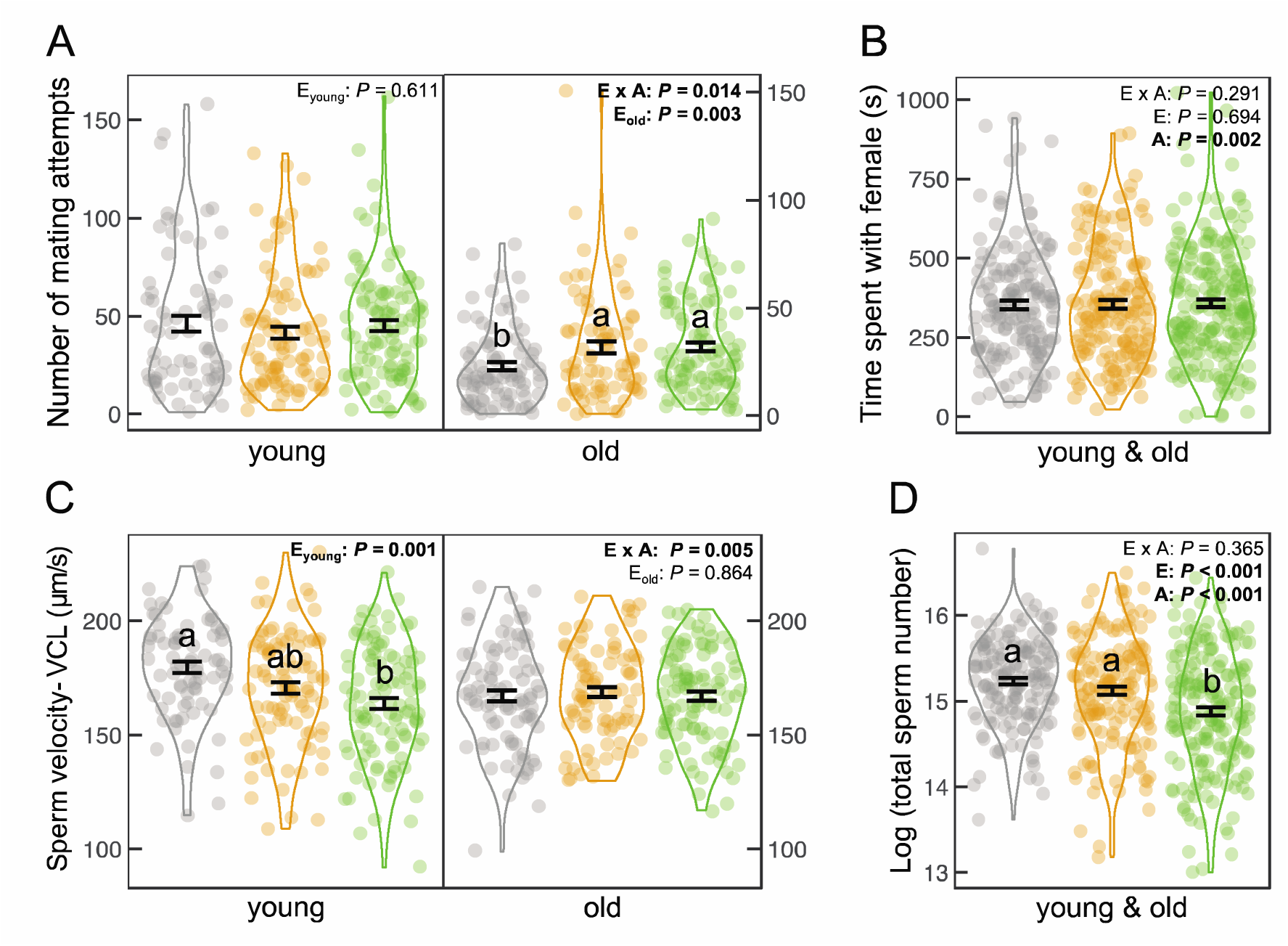
Effect of development environment and adult age on male reproductive effort. Colors indicate the environments: freshwater (F; grey); stable salinity (SS; orange); fluctuating salinity (FS; green). (A) number of mating attempts (*young* F: *n* = 75; SS: *n* = 92; FS: *n* = 107; *old* F: *n* = 89; S: *n* = 81; FS: *n* = 99), (B) time spent pursuing females (F: *n* = 168; SS: *n* = 181; FS: *n* = 210), (C) sperm velocity (*young* F: *n* = 77; SS: *n* = 94; FS: *n* = 105; *old* F: *n* = 90; SS: *n* = 85; FS: *n* = 102), (D) total sperm count (F: *n* = 162; SS: *n* = 167; FS: *n* = 196) are presented with the significance levels of environment (E), age (A) and their interaction (E×A) using bold font. Given a significant interaction, the environment effect was shown separately for each age class. Figures show mean ± *SE* and only indicate the conditional part of the results for (A) number of mating attempts. Different letters represent significant differences between environments based on Tukey’s tests (see Supplementary Table 5 for details).

The developmental environment affected the sperm velocity of young males (LMM, *F* = 7.014, *P =* 0.001), but not that of old males (LMM, *F* = 0.146, *P =* 0.864) (Fig. 3*C*). Sperm velocity was fastest for young males from the freshwater environment and slowest for males from the fluctuating salinity environment (pairwise: *P* < 0.001). Sperm from young males from the stable salinity environment had an intermediate velocity (both *P* > 0.103). The sperm velocity of young males depended on their body size (Supplementary Fig. 7). There was no environmental effect on sperm velocity after correcting for male size (Supplementary Table 7).

The developmental environment affected sperm count (LMM, *F* = 21.534, *P* < 0.001) irrespective of male age (i.e., no interaction: *F* = 1.014, *P* = 0.365) (Fig. 3*D*). Males from the fluctuating salinity environment produced fewer sperm than those from the other two environments (pairwise: both *P* < 0.001), with no significant difference between the latter two (*P* = 0.051). Controlling for male size, however, the sperm count was significantly lower for males from either saline environment than those from freshwater (pairwise: both *P* <0.018; Supplementary Table 7).

### Fluctuating salinity induced adult morphological changes

Environmental variability might cause developmental instability, creating maladaptive morphologies that lower fitness^59^. We found no effect of the developmental environment on skeletal deformity (i.e., spinal bending), but fluctuating salinity significantly affected the body shape of both sexes. It also reduced the length of the male intromittent organ^60^ (Supplementary Tables 8-11).

## DISCUSSION

Higher salinity has been shown to lower the fitness of freshwater organisms^32, 61, 62^, but it remains unclear whether fluctuations in salinity during development have more adverse effects than a consistently high salinity. We observed significant effects of both early-life higher salinity and its fluctuations on how fish allocate resources to a range of fitness-related traits. Fish that developed in a stable salinity environment reached maturity sooner, increased their reproductive effort and decreased their somatic maintenance. Fluctuations in salinity induced a different life-history response: slower development and lower investment in reproduction, with inconsistent changes in traits that facilitate somatic maintenance. Crucially, there were numerous sex-specific effects of fluctuations in salinity, including a mismatch in the timing of reproductive maturation between males and females.

Surprisingly, there was no differences in adult survival between fish reared in freshwater and the two saline environments. This is true, despite decreased investment in traits affecting somatic maintenance when fish developed in stable salinity. The absence of detectable survival costs alongside greater reproduction output makes it uncertain if there is a net fitness cost to stable salinity. In contrast, reduced reproductive investment when developing in fluctuating salinity should clearly lower fitness. Our results emphasize that environmental fluctuations – a common, but neglected phenomena in studies of freshwater salinization – can significantly lower reproduction and result in a greater decline in fitness than is expected when salinity is consistently high.

When developing in stable salinity rather than freshwater, fish grew faster, matured sooner, and invested more heavily in reproduction; but they also invested less into self-maintenance. Interestingly, the costs of stable salinity seemed to vary between the sexes: males from the stable saline environment had shorter relative telomere length (RTL), but there was no RTL differences for females. Rates of telomere shortening have been linked to the ability to cope with environmental challenges^63^. Faster growth and increased osmoregulation in a more saline environment are both associated with greater oxidative stress^64, 65^, which would account for accelerated telomere shortening^45, 66^. Why, however, was RTL shortening only seen in males? One explanation is a sex difference in metabolism, whereby females have more effective mechanisms than males to preserve energy, and their mitochondria exhibit greater resistance to oxidative stress^67^. As such, the energy-demanding processes of osmoregulation might result in greater energy consumption and higher oxidative damage in males. Another possibility is stronger selection for female longevity. Males in polyandrous species, like *G. holbrooki*, employ intensive ‘wear-and-tear’ reproductive strategies once they start to breed due to intense sexual selection^68, 69^. The price can be more rapid senescence due to greater investment in reproduction than maintenance. In contrast, female fecundity increases with both size and age (see *Results*), selecting for ongoing adult growth and greater somatic maintenance in females than males. This might also select for a better ability to cope with environmental stressors in females than males^70, 71^, affecting telomere shortening.

Nevertheless, shorter gut length in females from the saline than freshwater environment points to a ‘hidden’ cost of salinity. There is a well-documented trade-off between gut length and other energy-expensive tissues^53, 54^. It is plausible that reduced female gut length is an adaptive allocation strategy that prioritizes faster growth and maturation when developing in a stable salinity environment^72, 73^. Overall, our findings are broadly consistent with field observations of mosquitofish^61^, and similar patterns in sticklebacks^74^. These trends, often described as *r*-shifted strategies^74^ or a faster ‘pace of life’^17^, reveal that effects of stable salinity during development persist throughout adulthood.

When salinity was fluctuating rather than stable, there was no longer an effect on RTL in either sex, but females had a longer gut. Our findings suggest that fish experiencing frequent changes in their developmental environment might maintain somatic investment for future survival by allocating fewer resources to other processes. Indeed, fluctuations in salinity led to smaller eggs and a lower sperm count for young adults. Fluctuations in salinity also slowed juvenile growth, but the consequences were sex-specific. Females took longer to reach maturity but matured at the same size; while males took the same time to reach maturity but were therefore smaller upon maturation. Slower maturation delays the onset of breeding and is likely to lower fitness in a season breeder like *G. holbrooki*^75^. Our results corroborate studies in other fish species reporting slower development^76, 77^ and lower fecundity^78^ in more variable environments (albeit when temperature rather than salinity varied). Such damage had long-lasting effects. Old females carried fewer eggs and embryos, and old males had smaller sperm reserves even after they had been in the common garden freshwater environment for several months. In our study, most differences in reproductive traits between fish in a stable and fluctuating salinity environment disappeared when controlling for body size. This suggests that life-history shifts under salinity fluctuations are strongly driven by size-dependent reproductive output.

Our study provides insights into the impact of increased climatic variability on both the mean and variation in salinity within freshwater systems. It is assumed that developmental responses to elevated salinity impose fitness costs on freshwater fishes. In our study such costs were not readily apparent in the stable saline environment due to increases in reproductive traits likely to elevate fitness, with no detectable trade-off with survival. In contrast, costs of fluctuations in salinity were readily detected. Both males and females showed reductions in reproductive investment. We suggest that future studies pay greater attention to sex differences in the fitness consequences of fluctuations in stressors. While it is often suggested that males are more sensitive than females to environmental changes^67, 71, 79, 80^, our findings indicate that females can be more strongly affected by climate variability.

## METHODS

### Origin and maintenance of fish

We collected mosquitofish from a freshwater creek (35°180 27” S 149°070 27.9” E) in Canberra, Australia in 2020-2021. Adults were housed in 90L mixed-sex stock tanks filled with freshwater for 3 weeks. Pregnant females were transferred to individual 1L tanks to give birth (*n* = 118 broods). We randomly assigned a third of each brood to (a) freshwater control, (b) stable salinity or (c) fluctuating salinity treatment. This split-clutch design allowed us to control statistically for genetic variation and differences in maternal effects among broods that might affect trait expression. Siblings in the same treatment were housed in communal tanks (7 or 4L) at ≤ 1 individual/litre until they reached maturity.

Wild-caught juveniles were reared in mixed-sex stock aquaria until they could be sexed (a visible gravid spot for females and a pointed anal fin for males) and transferred to single-sex aquaria to ensure their virginity. Virgin females were used for male behavioural trials (see below).

All fish were kept under a 14 L: 10 D cycle at 28 ± 1°C and fed twice daily on either commercial fish flake or *Artemia* nauplii (for stock fish in aquaria) or *Artemia* nauplii only (fish in treatment tanks).

### Experimental overview

We set the level of salinity in treatment tanks by dissolving the appropriate amount of aquarium salt (Aquaforest, Brzesko, Poland) with tap water and aquatic conditioner (Seachem, Madison, GA, USA). Salinity levels were checked every two days using a refractometer. Juveniles in the stable salinity treatment were kept at a constant salinity level (10 ‰), while those in the fluctuating treatment were reared at salinities that cyclically increased and decreased cycle ever two days (10 ‰ → 20 ‰ → 10 ‰ → 0 ‰ → 10 ‰; mean = 10 ‰). We initially transferred newborn offspring assigned to the stable and fluctuating treatments into tanks with 2.5 ‰ saline water for two days as their first salinity exposure to allow for acclimation. No initial mortality occurred (personal observation).

Offspring assigned to the control treatment were placed in 0 ‰ water. Fluctuating salinity levels were created by manually rotating the fry between tanks of water with different salinity concentrations every two days. Juveniles in the control and stable treatments were also rotated among tanks with the appropriate salinity levels (0 or 10 ‰ respectively) to control for any effects associated with changing tanks.

We inspected treatment tanks every two days to check for sexual maturity and recorded the age of maturity. At this stage, all focal fish were still virgins. To minimize immediate physiological effects of the juvenile environment^81^, we transferred newly matured adults into separate 1L freshwater tanks for 1 week before making any measurements. We recorded their standard length (SL: the snout tip to the base of caudal fin), gonopodium length and body shape. Females (*n* = 290) and males (*n* = 240) were randomly selected (at least one male and one female per brood/treatment) and euthanized to determine a suite of early life-history traits (immune response, relative gut length, relative telomere length, egg number, egg size).

To test for longer-term effects of the developmental environment, we introduced the remaining adults (at least one male and one female per brood/treatment) into mating tanks. We individually housed each focal female (*n* = 316) or male (*n* = 327) with a wild-caught fish of the opposite sex in 4L freshwater tanks for 12 weeks (±75% of the breeding season length in the source population^75^). Focal adults were free to mate and bred with the stimulus fish.

Before being placed into mating tanks, male mating behavior and sperm traits were measured (see below). The wild-caught stimulus fish were rotated fortnightly to maintain the sexual interest of the focal individuals. Any deaths were recorded to test for an effect of the developmental environment on adult survival. After 12-weeks, we again measured the life-history and reproductive traits of the surviving females (*n* = 284 of 316; 90%) and males (*n* = 277 of 327; 85%).

We tested the following traits at: (a) 1 week after maturation (young); (b) after another 12 weeks (old).

### Body length, body shape and gonopodium length

Focal fish were anaesthetized in ice-cold water and placed on a glass slide. The side of their body was photographed to measure their SL and gonopodium length using *ImageJ*^82^. We also measured their body shape by digitizing 10 landmarks using tpsDig (see Booksmythe et al^60^ for our protocol). Any skeletal deformities were recorded. Spinal bending occurs at a low rate in the wild (personal observation).

### Growth

We placed a juvenile (maximum five fry per brood/treatment; *n* = 888) in a shallow container containing some of their tank water and photographed it from above fortnightly (0, 2, 4 and 6 weeks after birth) to determine its SL. We excluded week 8 as, by then, some juveniles had started to mature (Fig. 1*A*). Fry are too small to be individually marked, so we used the mean value per brood.

To measure adult growth, we followed the same procedure but photographed *every* adult that went through the mating period (0, 2, 4, 6 and 8 weeks after their initial week as an adult in freshwater).

### Male mating behavior

We placed each male into a 7L tank with a randomly selected virgin female (SL: 29.8 ± 0.09 mm; *n* = 559) who was behind a mesh screen. After 10 minutes, we removed the screen and recorded the following male behaviors for 20 minutes: (a) number of mating attempts (the gonopodium swings forward under the female’s gonopore), (b) number of successful mating attempts (the gonopodium contacts the gonopore^57^), (c) time in association with the female (distance <1 SL of the male).

### Total sperm count

After the behavioral trial, the male was anaesthetized in icy water, put on a glass slide covered with a 1% polyvinyl alcohol solution and then placed under a dissecting microscope. His gonopodium was swung forward, and we gently pressed on the abdomen to empty his sperm reserves. Males were then returned to their individual tanks for 7 days to allow for sperm replenishment^83^. We examined the count and velocity of replenished sperm to standardize sperm age and to avoid any effects on sperm count of differences in copulation rates during the mating trial. After 7 days, the males were re-stripped to collect the sperm into a known volume of extender medium (207 mM NaCl, 5.4 mM KCl, 1.3 mM CaCl2, 0.49 mM MgCl2, 0.41 mM MgSO4, 10 mM Tris (Cl); pH 7.5) using a 100-μl pipette. The sperm solution was subsequently vortexed for 30 seconds and mixed using 10-ul pipette several times. We placed 3μl of the solution on a 20-micron capillary slide (Leja) to measure sperm number using the program CEROS Sperm Tracker (Hamilton Thorne Research, Beverly, MA, USA) under 100× magnification. The sperm number was analyzed using the mean value of five randomly selected subsamples per male (repeatability: *r* ± SE = 0.829 ± 0.010, *P* < 0.001, *n* = 525 male-ages). The count data of 28 samples were not tested due to the unavailability of slides during the Covid-19 lockdown.

### Sperm velocity

While stripping sperm reserves from focal males under the microscope, we collected two samples (each of 4 sperm bundles) into individual Eppendorf tubes containing 2μl of extender medium. Sperm velocity was calculated as the weighted average of the motile sperm tracks in both samples. In each Eppendorf tube, we pipetted the sperm solution and placed 3μl into the centre of a cell in a 12-cell multi-test slide (MP Biomedicals, USA) covered with 1% polyvinyl alcohol solution. We activated the sperm sample using a 3μl solution of 125 mM KCl and 2 mg/ml bovine serum albumin for 30 seconds and covered it with a coverslip. Using the CEROS Sperm Tracker (Hamilton Thorne Research), we measured (a) average path velocity (VAP): the average velocity over a smoothed cell path, (b) curvilinear velocity (VCL): the actual velocity along the trajectory, and (c) straight-line velocity (VSL). Because VAP and VSL were highly correlated with VCL (VAP-VCL: *r* = 0.996; VSL-VCL: *r* = 0.998, *n* = 553), we focused on the actual velocity of sperm cells (VCL)^58^.

### Female fecundity

After immune responses were measured (see below), females were euthanized to measure their fecundity. For young females (i.e., still virgins), we counted and photographed their unfertilized, mature eggs and measured the diameter of each egg using *ImageJ*. For old females (i.e., post the mating period), we measured the number of: (a) embryos (i.e., fertilized eggs); (b) unfertilized eggs; and (c) offspring born during the 12-week mating period. We also noted how many times females produced a brood of offspring during the 12 weeks (range: 0 – 2). We defined (a) + (c) as the total number of offspring; and (a) + (b) as the total number of eggs for old females.

### Immune response

We used a phytohemagglutinin (PHA) injection assay to quantify the cell-mediated immunity of mosquitofish^84, 85^. When fish were under anaesthesia, we recorded the thickness of their body at the posterior end of the dorsal fin with a pressure-sensitive spessimeter (Mitutoyo 547- 301; accuracy: 0.01 mm; average of five measurements per fish). At the point where the body thickness had been measured, we then injected 0.01mg of PHA dissolved in 0.01ml of PBS into the left side of the fish. After 24 hours, the body thickness was again measured five times (repeatability: *r* ± SE = 0.992 ± 0.000, *P* < 0.001, *n* = 1114 measure-female-age; 0.978 ± 0.001, *P* < 0.001, *n* = 1022 measure-male-age). We treated the difference in mean thickness before and after the injection as an index of the immune response.

### Gut length

To test if gut length was affected by salinity and its fluctuations during development, we euthanized males and females after the PHA assay. Their guts were dissected under a microscope then photographed. Gut length was measured with *ImageJ*.

### Relative telomere length (RTL)

We collected the tail muscle after removing the attached skin and skeleton and then temporarily stored it at –80°C in 70% ethanol. For each sex at each age, we randomly selected 30 broods that had at least one individual per treatment to control for telomere variation across broods. Genomic DNA was extracted using DNeasy Blood and Tissue Kit (QIAGEN), and its concentration was quantified using Qubit™ dsDNA BR Assay Kits (Thermo Fisher Scientific Inc.), then stored at –20 °C. Using real-time qPCR, we measured RTL: how the sample differed from a reference DNA sample in term of the ratio (T/S) of telomere repeat copy number (T) divided by a single-copy gene copy number (S). Following a previous *Gambusia* study^86^, Tel1b (5’- CGGTTTGTTTGGGTTTGGGTTTGGGTTTGGGTTTGGGTT-3’) and Tel2b (5’-GGCTTGCCTTACCCTTACCCTTACCCTTACCCTTACCCT-3’) were used as the telomere primers. The control single-copy gene Melanocortin 1 receptor (MC1R) was amplified using primers MC1R-F (5’-CCTGTAGGCGTAGATGAGCG-3’) and MC1R-R (5’-CACCAGTCCCTTCTGCAACT-3’). We confirmed that the primers bind to a single region (Oliver Stuart, personal observation).

For each sample of telomere and MC1R, reactions were run in triplicate on 96-well plates using the QuantStudio^®^ 3 qPCR system. Each well contained 1ul of 20ng/ul DNA sample with 9ul of master mix, including 5ul 2x SensiMix SYBR No-ROX (Meridian Bioscience Inc), 3.4ul Milli-Q water, 0.3ul of 10uM for both forward and reverse primers. For telomere amplification, an initial step started at 95°C for 10 minutes following by a total of 40 cycles of 95°C for 15 seconds, 60°C for 15 seconds and 72°C for 15 seconds. Afterwards, a melt curve stage (95°C for 15 seconds, 60°C for 60 seconds and 95°C for 15 seconds) was created to ensure qPCR specificity. For MC1R amplification, we established a 3-minute denaturation at 95°C, then a total of 40 PCR cycles at 95°C for 15 seconds, 60°C for 30 seconds and 72°C for 20 seconds, and a melt curve analysis (95°C for 15 seconds, 60°C for 60 seconds and 95°C for 15 seconds) after each run. Standard curves for both telomere and MC1R were generated using five serial dilutions of DNA (0.01, 0.1, 1, 5 and 20 ng/µL) run on each plate with an acceptance threshold (100 ± 20 %) of amplification efficiency (E), using the formula E = 10 ^(-1/slope of the standard curve)^. A negative control and two inter-plate calibrators (i.e., reference samples) were also run on each plate. RTL was assessed by the mean crossing threshold (Ct) values of each sample using the following equation^87^:

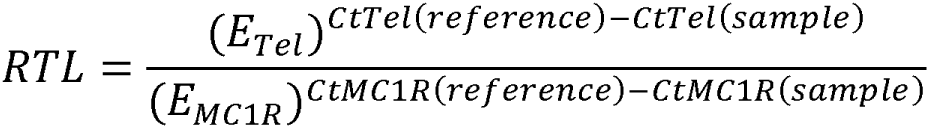

*E* is the mean amplification efficiency of either telomere or MC1R on each plate. *CtTel(reference)* and *CtMC1R(reference)* are the mean Ct values of the two calibrator DNA samples in the plate for telomere and MC1R, respectively. *CtTel(sample)* and *CtMC1R(sample)* are the average Ct values from the three repeated measurements of each sample for telomere and MC1R, respectively. In total, 29 of 2880 reactions (∼1%) were excluded as undetermined data due to pipetting error. The Ct values of each sample were highly repeatable (MC1R: *r* ± SE = 0.980 ± 0.002, *P* < 0.001; telomere: *r* ± SE = 0.950 ± 0.004, *P* < 0.001, *n* = 465).

### Statistical analyses

Our analyses were pre-registered online (<u>osf.io/8s4h3</u>). The significance level was set at α = 0.05 (two-tailed). Analyses were run using R v4.0.5 in R studio v1.3.1093. We used the *Anova* function of *car* package (type III Wald *chi-square* or *F*-tests with Kenward-Roger *df*) to determine *P* values. All summary statistics are presented as mean ± SE.

### All traits (except growth, mortality and morphology)

*Analysis 1 (main effects of the developmental environment and adult age):* To test the effect of the developmental environment on measured traits and whether it was age-dependent, we considered environment type (freshwater, stable salinity, fluctuating salinity), age (young, old) and their interaction as fixed factors in initial models. Age was excluded if a trait was only measured at one age (e.g., egg size). If there was a significant interaction, we examined the environment effect separately at each age to better understand the phenotypic change at different life stages. If the interaction was non-significant, it was removed from the model to estimate the main effects of environment and age^88^. The best fit of the model remained unchanged after excluding the non-significant interaction (Supplementary Information).

Given a significant environment effect, we ran post-hoc Tukey’s tests to test the significance of the three possible pairwise differences. Each measured trait was analyzed with a separate model. All models were run separately for each sex because there are well-known sex differences in life history traits in *G. holbrooki* ^60, 75, 89, 90^. Brood identity (ID) was included in all models as a random factor using (1 + age | brood ID) to account for differences among broods in mean values and any difference in the effect of age. We also included individual ID as a random factor for traits that were repeatedly measured when young and old (namely, body deformity and male reproductive traits).

For most traits, we ran linear mixed models (LMMs) because the model residuals fulfilled the assumption of normality, homoscedascity and linearity as determined via Q-Q plots. For other traits, we either log-transformed the data (sperm count, gut length) or ran generalized linear mixed models (GLMMs) with the appropriate error distribution: (a) quasi-Poisson error for total egg number, (b) negative binomial error for total egg number of young females and immune response, and (c) negative binomial error with zero-inflation for the number of mating attempts, total number of offspring and number of embryos. We included the same variables in zero-inflated models as those in main models. For traits that contained mostly, or only, 0 and 1 responses, we set the >1 data to 1 and ran models using (d) binomial error, including male successful copulation (only 4 % of the males copulated successfully more than once), female brood number (only 2 % of females had more than 1 brood) and body deformity (bending or not). Due to convergence issue, we used (1| brood ID) instead of (1+ age | brood ID) when testing the environment-by-age interaction in the initial models for immune response, number of mating attempts and body deformity, as well as in the reduced model (i.e., interaction removed) for the main effects of age and environment on deformity. We also removed (1| brood ID) in the zero-inflated model of the number of mating attempts by old males. For log-transformed gut length, we accounted for its allometric relationship with body size^91^ by log-transforming absolute SL and including the transformed SL as a covariate (standardized) to test for treatment effects on relative gut length. For RTL, we ran an additional LMM including age since birth (standardized) as a covariate (with main effects of developmental environment and adult age) to test whether individuals with a longer juvenile period (i.e., older absolute age) had a shorter RTL. Finally, we ran overdispersion tests using the *DHARMa* package to ensure that the variance was not greater than the mean for GLMMs.

### Analysis 2 (controlling for body size)

A significant effect of the developmental environment and/or adult age for some traits might be due to their effect on body size (i.e., intermediate outcome^91^), especially when body size differed among environments or increased with age (Fig. 1). To test if body size influenced the measured traits, we ran two additional analyses. First, we included the absolute SL (globally standardised across all treatments: mean = 0, SD = 1) as a covariate in the models described in *Analysis 1*. If the inclusion of absolute SL changed the significance of the main effects of environment or age, this implies that these effects were largely attributable to allometric effects driven by size differences among environments or changes in size with age. Second, it is also possible that relative body size within each treatment affects traits. To assess this, we ran models including SL standardized within each treatment/age combination (mean = 0, SD = 1) as a covariate^91^. For gonopodium length, we log-transformed SL before standardizing it, as well as log-transforming gonopodium length^91^.

### Analysis of growth and mortality

We ran a GLMM with Poisson error for juvenile growth and separate LMMs for adult growth for each sex. We considered SL as the response variable, and treated environment as a fixed factor, and age (in weeks) as a fixed covariate. We included the environment-by-age interaction, as well as age^2^ to account for non-linear growth. We treated brood ID (for both juveniles and adults) and individual ID (for adults only) as random factors to control for repeated measures from the same family and/or fish. If the environment-by-age interaction was significant, we ran individual LMMs (and post-hoc pairwise tests) at each age to determine when body size started to diverge significantly between environments.

Adult mortality was evaluated separately for each sex using survival analysis. We ran Cox regression models (*coxme* package) with environment as the fix factor and brood ID as a random factor. Fish still alive at the end of the experiment were treated as right-censored data.

### Analysis of morphology

Body shape was analysed using Procrustes multivariate analysis of variance (*geomorph* package) with 10 landmarks to describe body shape^60^. In brief, body size was scaled to an average size (centroid size of 1), and the morphometric analysis provided Procrustes coordinates and centroid size for each sample. We then ran separate LMMs for each sex with the two-dimensional set of coordinates as the response variable, and environment, age and their interaction as predictors, with log-transformed centroid size as a covariate and brood ID as a random factor. If the environment-by-age interaction was non-significant, it was excluded from the final model. Significant differences between groups were tested for using post-hoc pairwise comparisons of Procrustes distances. We excluded fish with spinal deformities (as the effect of environment and age on skeletal deformity was addressed in *Analysis 1*). To test for an age effect, we only used individuals that were measured when both young and old to better control for individual variation (females *n* = 252; males *n* = 247).

## Data and code availability

Data generated and analyzed in this study and all codes have been deposited at Dryad (https://datadryad.org/stash/share/wwDDw71Wd2TZw4J0B3kMdiGFzbq5vrkK_SG_IlyfC0k) and will be publicly available as of the date of publication.

## Acknowledgements

We thank U. Aich for help with sperm analysis; O. Stuart, M. Ruibal and N. Aitken for help with telomere measurement; M. Head and P. Tiatragul for help with shape analysis; the staff of ANU Animal Services for help with fish maintenance. M.D.J. and D.W.A.N. were supported by Australian Research Council Discovery Project grants (DP190100279: M.D.J.; DP210101152 : D.W.A.N.). M.-H.J.C was supported by an Australian Government Research Training Program International Scholarship and a Taiwanese Government Scholarship for Study Abroad.

## Author contributions

M.-H.J.C., R.J.F., and M.D.J. conceived the study. M.-H.J.C., D.W.A.N., R.J.F., and M.D.J. designed the experiment. M.-H.J.C. and L.M.H. developed the methodology of telomere measurement. M.-H.J.C. collected all data, performed the analyses, visualized the data, and interpreted the results. D.W.A.N. and M.D.J contributed to data analysis and interpretation. M.-H.J.C. wrote the first draft, with D.W.A.N. and M.D.J. providing critical feedback and revisions.

## Additional information

Supplementary information is provided as a separate file for the reviewing process.

