## Supplementary Information for "Fluctuating salinity during development impacts fish productivity"

Supplementary figures

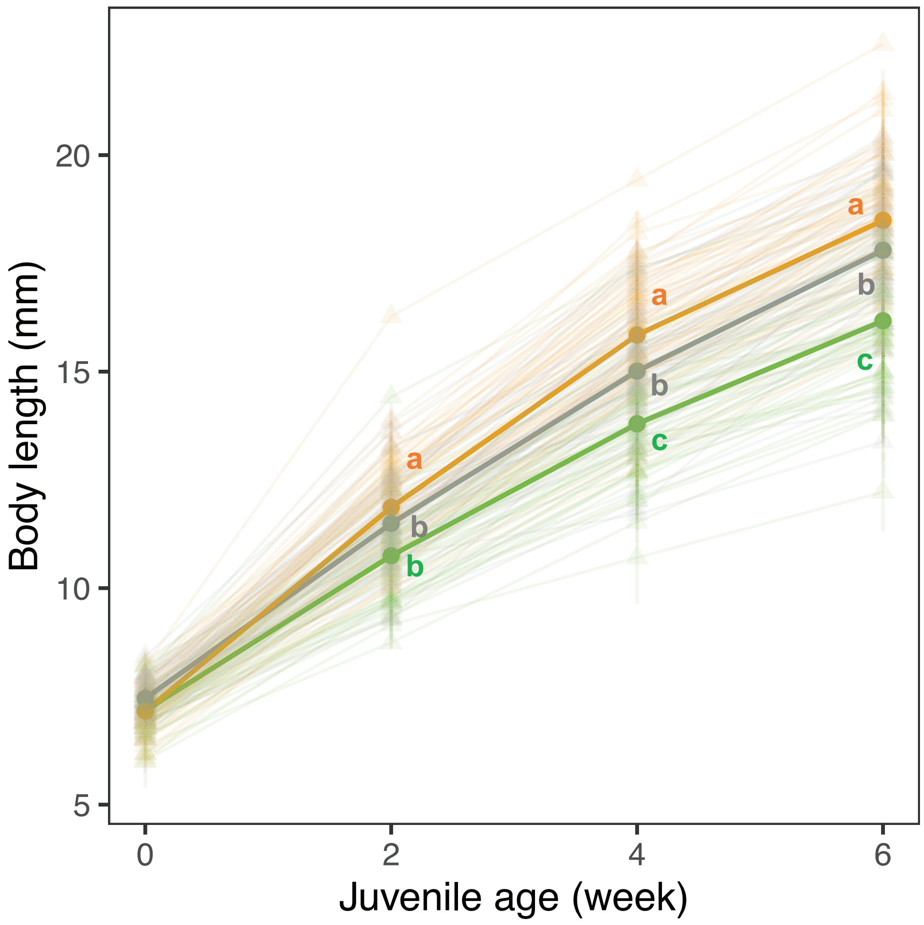


**Figure S1. Effects of salinity and its fluctuation level on juvenile growth.** Colors indicate the environment: freshwater (grey); stable salinity (orange); fluctuating salinity (green). Changes in juvenile length ($n$ = 888 with 3282 observations) are shown at the family level ($n$ = 91 broods) (triangle, thin line) along with the mean for each environment (circle, thick line). Different letters represent significant differences between environments from Tukey's tests. Line bars represent mean ± *SE*.

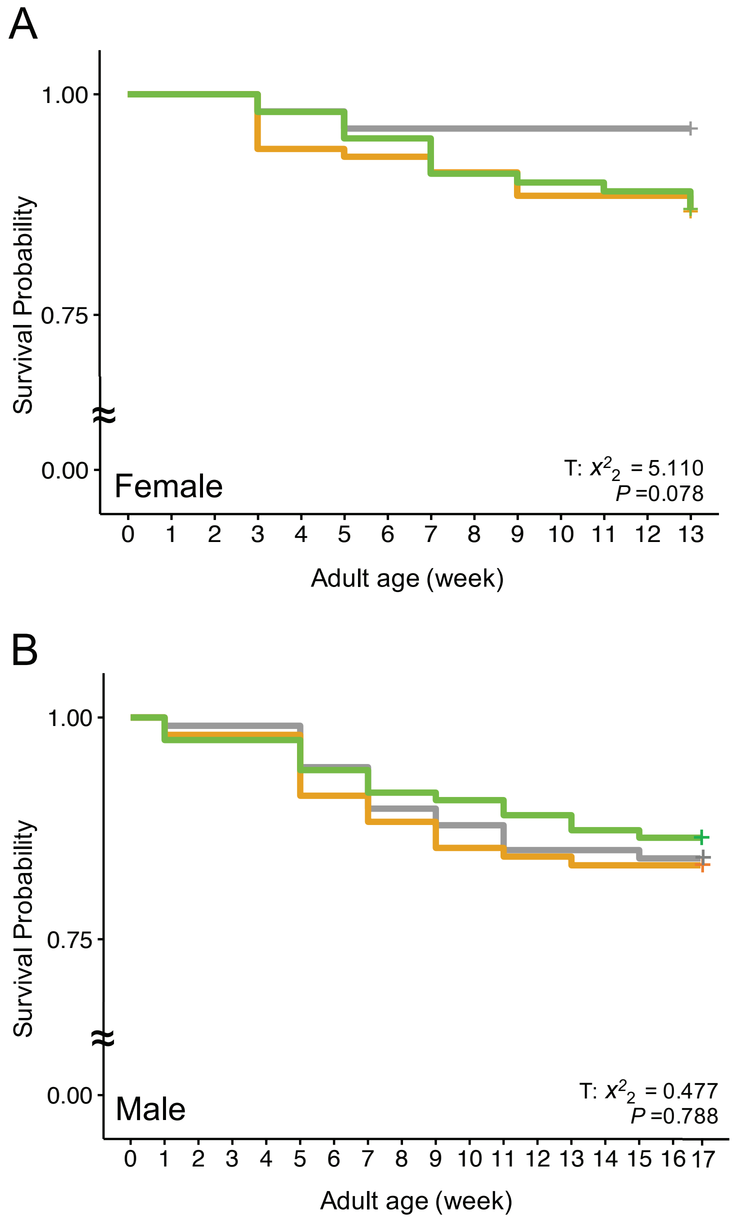


**Figure S2. Effects of developmental environment on adult mortality.** Environments (E) are shown using colors: freshwater (grey; female: $n$ = 103; male: $n$ = 107), stable salinity (orange; female: $n$ = 113; male $n$ = 102) and fluctuating salinity (green; female: $n$ = 100; male: $n$ = 118) with test statistic and $P$ value (see Table S4*A* and S5*A* for details).

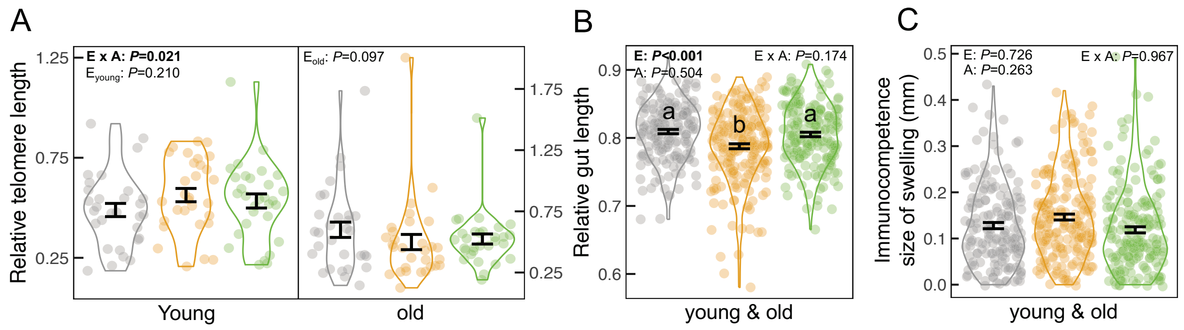


**Figure S3. Effects of developmental environment and adult age on female traits linked to somatic maintenance.** Colors indicate the environments: freshwater (F; grey); stable salinity (SS; orange); fluctuating salinity (FS; green). (A) relative telomere length (*n* = 30 per treatment/age) (B) relative gut length (F: *n* = 171; SS: *n* = 213; FS: *n* = 186), (C) immune response (F: *n* = 168; SS: *n* = 205; FS: *n* = 184) are shown along with the significance levels of environment (E), age (A) and their interaction (E×A) using bold font. If there was a significant interaction, the environment effects are shown separately for each age class. Error bars indicate mean ± *SE*. Different letters represent significant differences between environments from Tukey's tests.

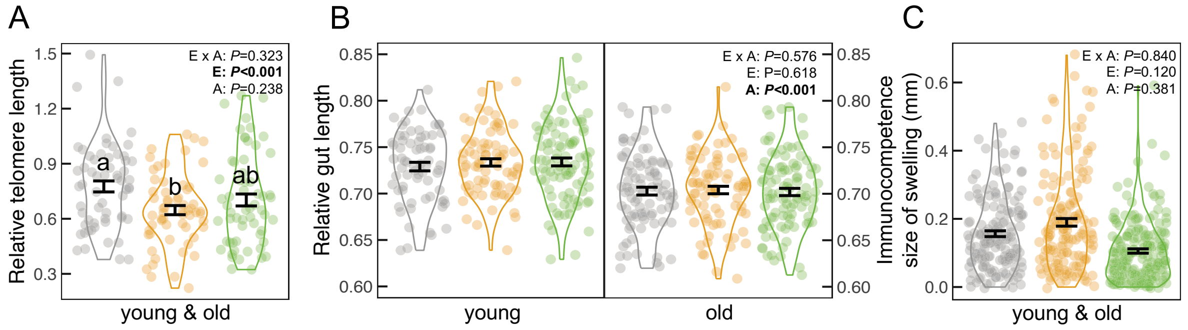


**Figure S4. Effects of developmental environment and adult age on male traits linked to somatic maintenance.** Colors indicate the environments: freshwater (F; grey); stable salinity (SS; orange); fluctuating salinity (FS; green). (A) relative telomere length (*n* = 30 per treatment/age), (B) relative gut length (*young* F: *n* = 68; SS: *n* = 79; FS: *n* = 93; *old* F: *n* = 90; SS: *n* = 85; FS: *n* = 102), (C) immune response (F: *n* = 157; SS: *n* = 163; FS: *n* = 191) are shown with the significance levels of environment (E), age (A), and their interaction (E×A) in bold font. If there was a significant age effect, the environment effects are shown separately for each age class. Different letters represent significant differences between environments from Tukey's tests. Error bars indicate mean ± *SE*.

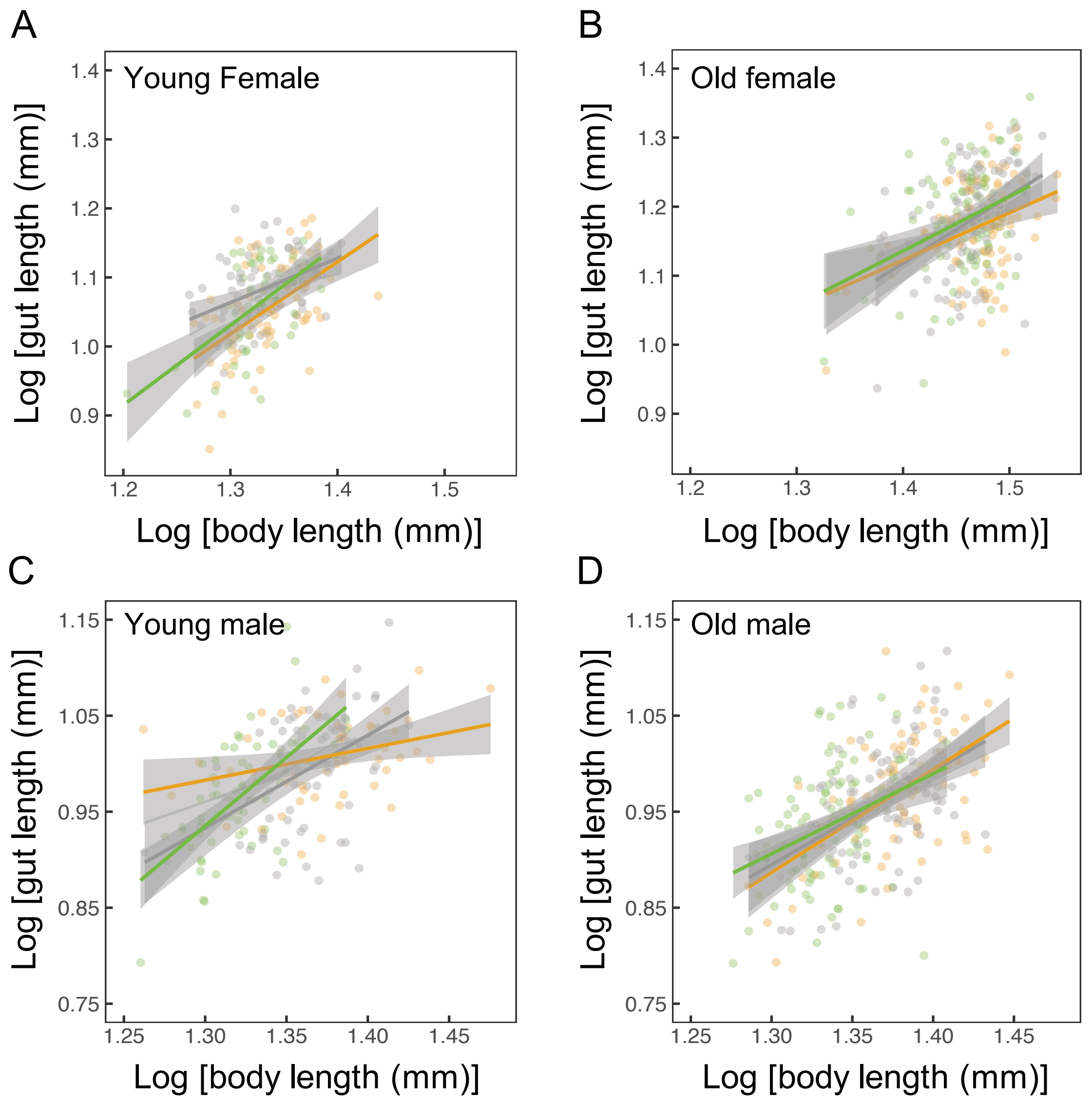


**Figure S5. Relationship between body size (log-transformed) and gut length (log-transformed).** (A) Young female; (B) old female; (C) young male; and (D) old male. Colors represent three developmental environments: freshwater (grey); stable salinity (orange); fluctuating salinity (green). The allometric relationships are shown along with regression lines and 95% confidence intervals. The statistical outputs are in Table S4*C* and S5*C*.

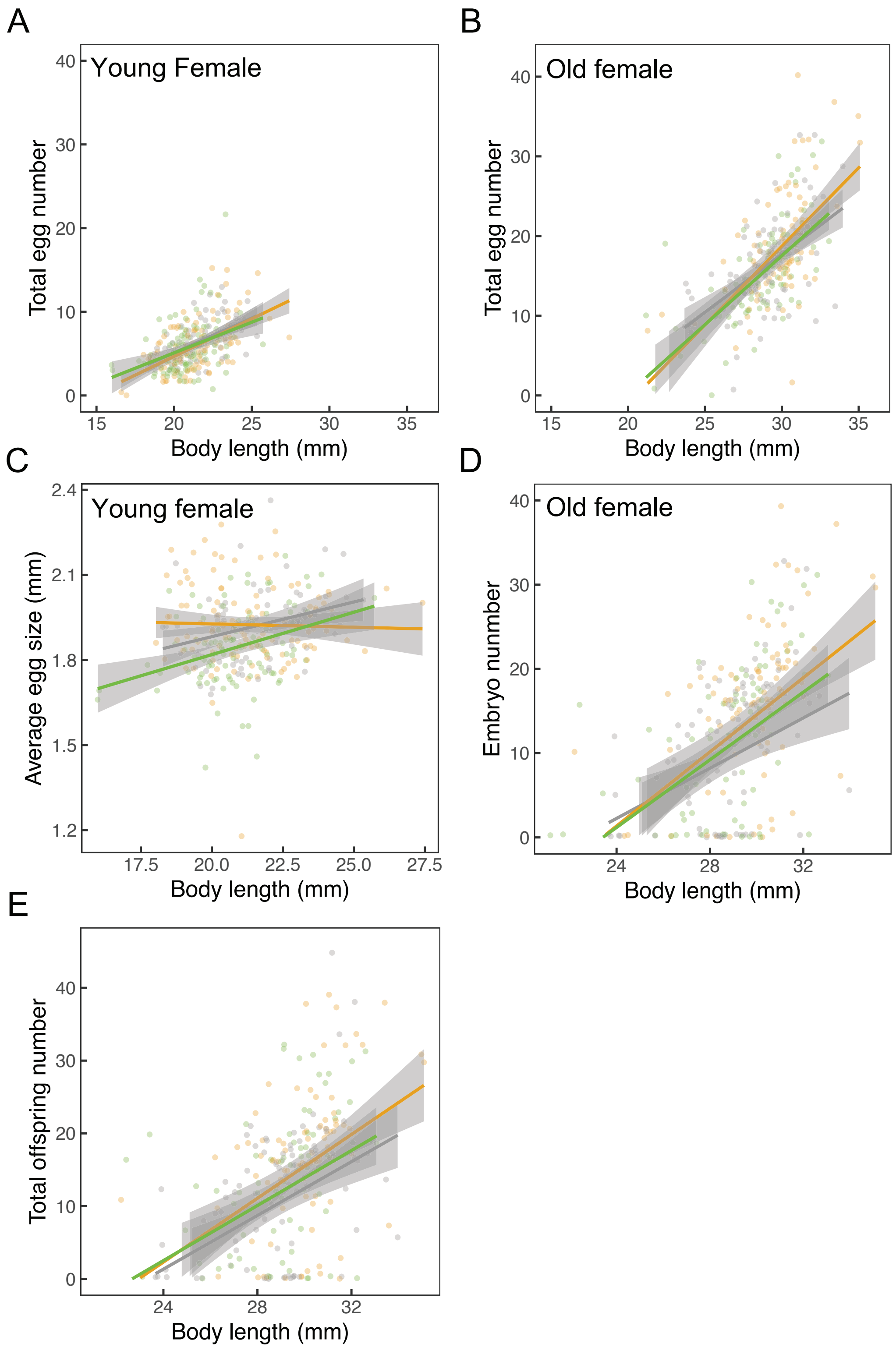


**Figure S6. Relationship between female fecundity and body size.** (A) Total egg number of young females; (B) total egg number of old females; (C) average egg size of young female; (D) embryo number of old female; and (E) total offspring number. Colors represent three developmental environments: freshwater (grey); stable salinity (orange); fluctuating salinity (green). The statistical outputs are shown in Tables S6*C*-*G*. The allometric relationships are shown along with regression lines and 95% confidence intervals.

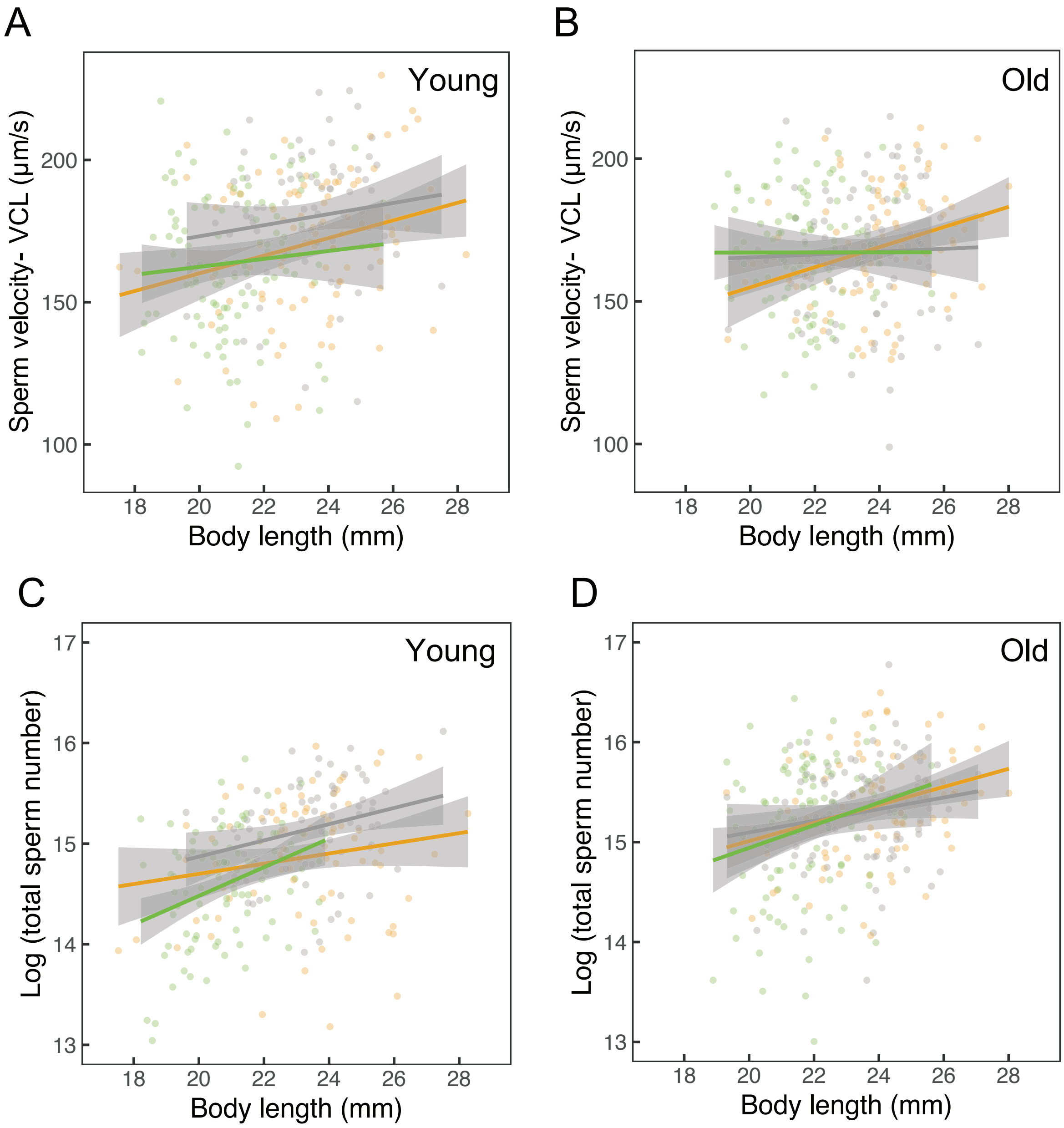


**Figure S7. Relationship between sperm traits and male body size.** (A) Sperm velocity of young males; (B) sperm velocity of old males; (C) log-transformed sperm number of young males; and (D) log-transformed sperm number of old males. Colors represent three developmental environments: freshwater (grey); stable salinity (orange); fluctuating salinity (green). The statistical outputs are shown in Tables S7*F* and S7*G*. The allometric relationships are shown along with regression lines and 95% confidence intervals.

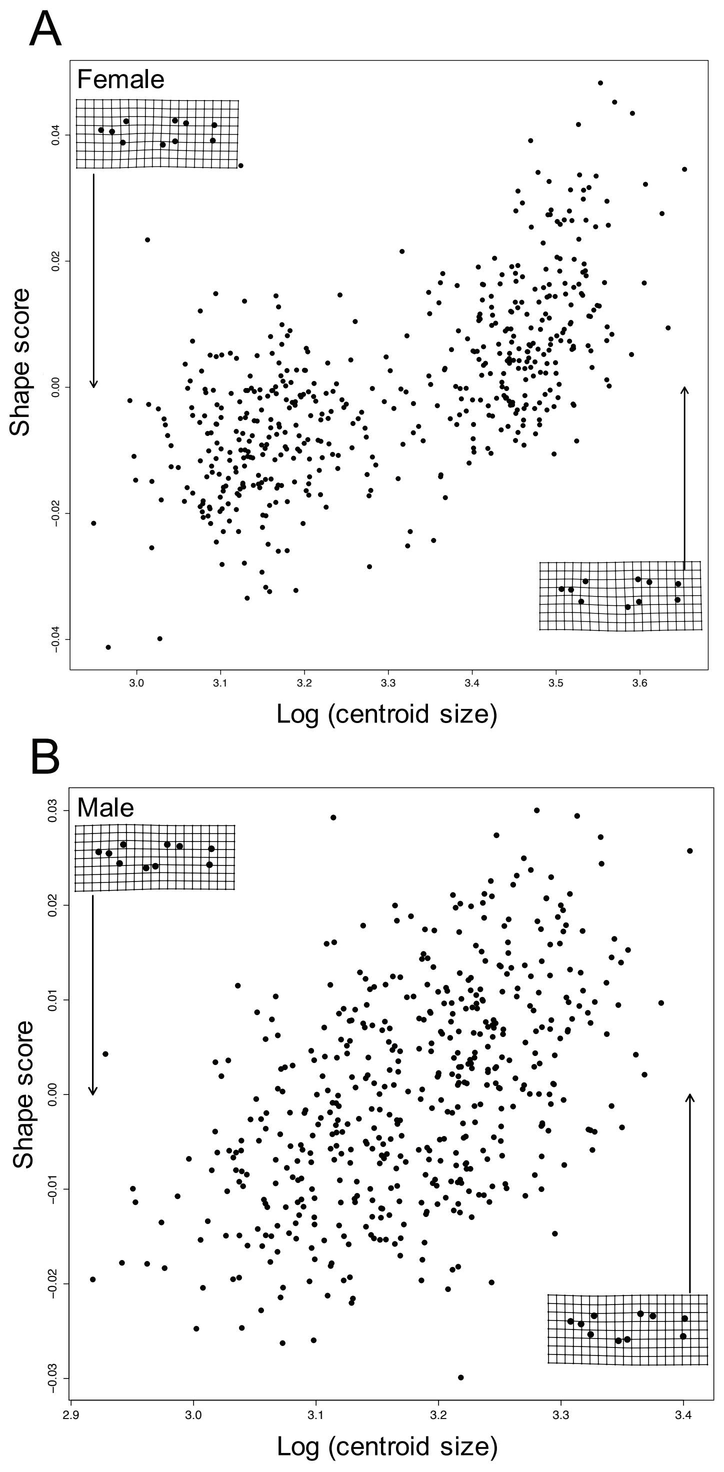


**Figure S8. Relationships between body size and shape of (A) females and (B) males.**

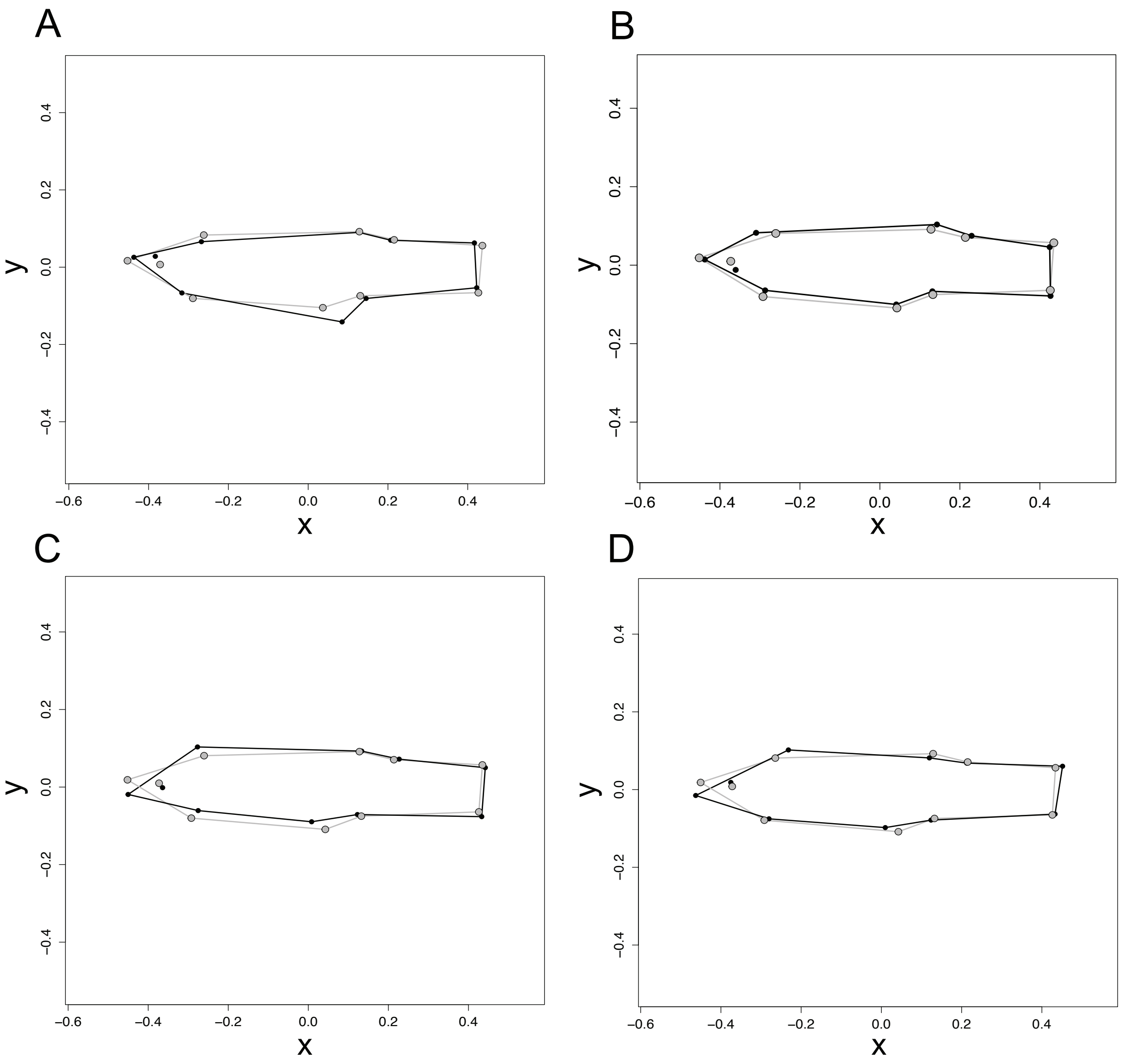


**Figure S9. Mean female body shape for each developmental environment and adult age.** The shape of: (A) young (grey) and old (black) females; (B) freshwater (grey) and stable-salinity (black) females; (C) freshwater (grey) and fluctuating-salinity (black) females; (D) stable-salinity (grey) and fluctuating-salinity (black) females. Female shape differences are magnified by 5x for the age comparison and 12x for the treatment comparisons to aid visualisation. X and y axes represent standardized shape co-ordinates.

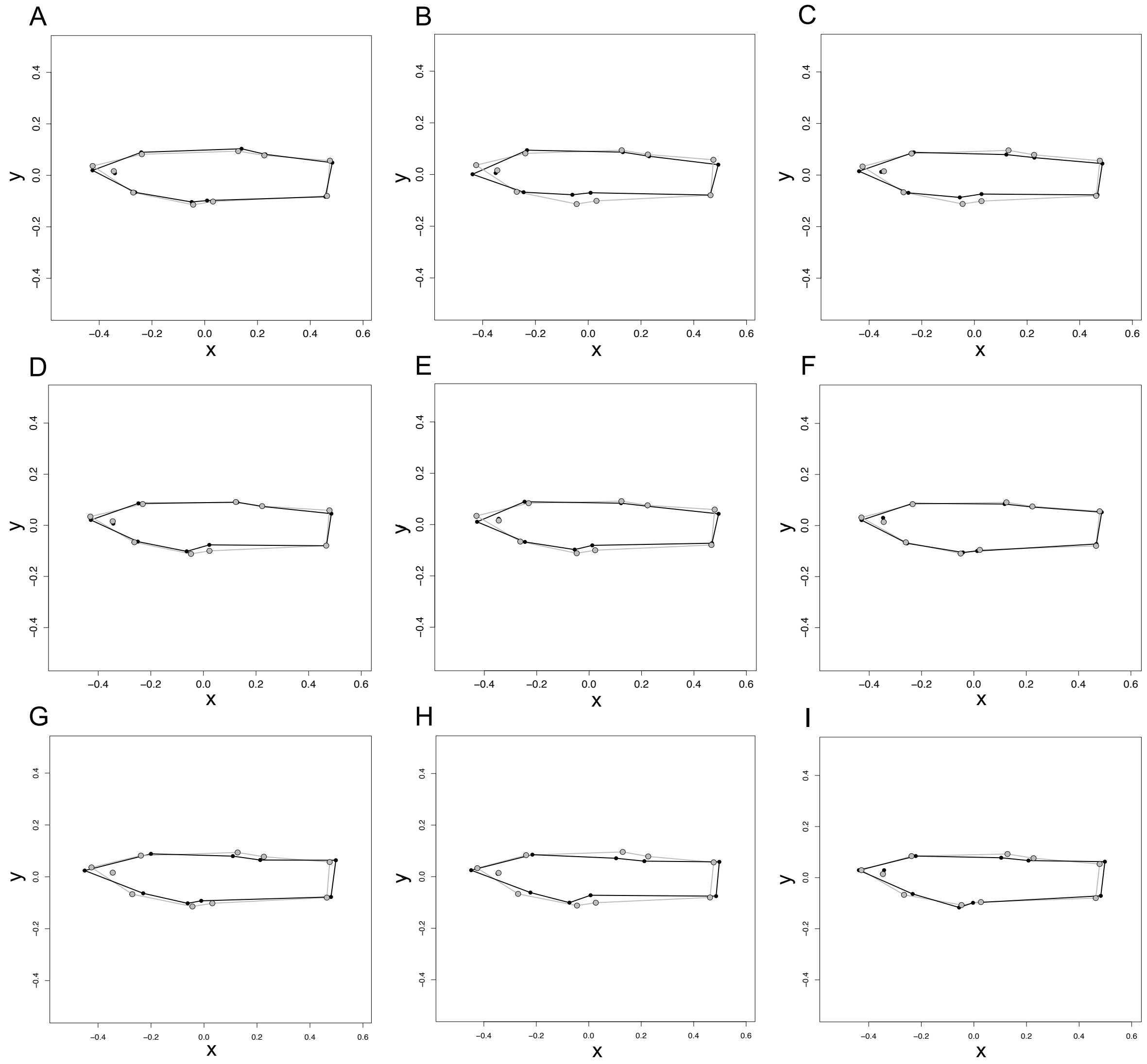


**Figure S10. Mean male body shape for each developmental environment and adult age**. The shape of young males: (A) freshwater (grey) and stable-salinity (black) males; (B) freshwater (grey) and fluctuating-salinity (black) males; (C) stable-salinity (grey) and fluctuating-salinity (black) males. The shape of old males: (D) freshwater (grey) and stable-salinity (black) males; (E) freshwater (grey) and fluctuating-salinity (black) males; (F) stable-salinity (grey) and fluctuating-salinity (black) males. The shape of young (grey) and old (black) males in (G) freshwater; (H) stable salinity; (I) fluctuating salinity environment. Male shape differences are magnified 5x to aid visualisation. X and y axes represent standardized shape co-ordinates.

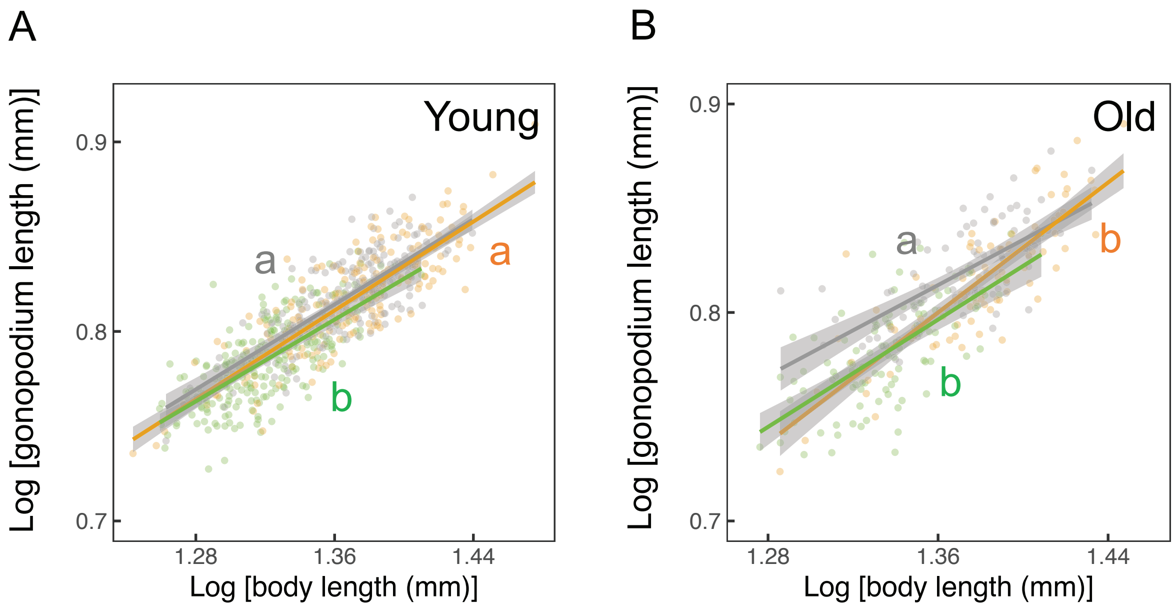


**Figure S11. The allometry of gonopodium length relative to absolute body length of young and old males.** Colors represent three environments: freshwater (grey) (young: $n$ = 190; old: $n$ = 90); stable salinity (orange) (young: $n$ = 207; old: $n$ = 85); fluctuating salinity (green) (young: $n$ = 231; old: $n$ = 102). Different letters indicate significant differences between environments (Table S17). Error bars represent mean ± *SE.* The allometric relationships are shown along with regression lines and 95% confidence intervals.

Supplementary tables

**Table S1. Statistical outputs of environment effect on growth of (A) juveniles, (B) adult females and (C) adult males.**

### (A) Juvenile growth

(i) Model with environment*age interactive effect

| **Fixed effect** | Estimate | *SE* | $\chi^2$ | *df* | *P* |
|---|---|---|---|---|---|
| Intercept (freshwater) | 1.989 | 0.019 | 11217.480 | 1 | **<0.001** |
| Environment (fluctuating) | -0.034 | 0.024 | 3.059 | 2 | 0.217 |
| Environment (stable) | 0.003 | 0.024 | | | |
| Age | 0.031 | 0.009 | 12.291 | 1 | **<0.001** |
| Age$^2$ | 0.286 | 0.021 | 187.671 | 1 | **<0.001** |
| Environment (fluctuating) * Age | -0.009 | 0.006 | 12.281 | 2 | **0.002** |
| Environment (stable) * Age | 0.010 | 0.006 | | | |

| **Random effect** | Variance | *sd* | Number of groups |
|---|---|---|---|
| Brood ID | 0.001 | 0.032 | 91 |

Due to a significant environment*age interaction, environment effects were tested for separately at each age:

(ii) Juvenile age: Week 0

| **Fixed effect** | Estimate | *SE* | *F* | *df* | *df.res* | *P* |
|---|---|---|---|---|---|---|
| Intercept (freshwater) | 7.301 | 0.050 | 21407.719 | 1 | 117.030 | **<0.001** |
| Environment (fluctuating) | -0.051 | 0.034 | 1.986 | 2 | 834.140 | 0.138 |
| Environment (stable) | -0.067 | 0.034 | | | | |

| **Random effect** | Variance | *sd* | Number of groups |
|---|---|---|---|
| Brood ID | 0.182 | 0.427 | 91 |
| Residual | 0.120 | 0.347 | |

(iii) Juvenile age: Week 2

| **Fixed effect** | Estimate | *SE* | *F* | *df* | *df.res* | *P* |
|---|---|---|---|---|---|---|
| Intercept (freshwater) | 11.086 | 0.113 | 9641.140 | 1 | 117.350 | **<0.001** |
| Environment (fluctuating) | -0.035 | 0.074 | 202.730 | 2 | 775.820 | **<0.001** |
| Environment (stable) | 1.093 | 0.074 | | | | |

| **Random effect** | Variance | *sd* | Number of groups |
|---|---|---|---|
| Brood ID | 0.933 | 0.966 | 91 |
| Residual | 0.520 | 0.721 | |

| **Pairwise comparison** | *P* |
|---|---|
| Freshwater-Fluctuating | 0.887 |
| Freshwater-Stable | **<0.001** |
| Stable-Fluctuating | **<0.001** |

(iv) Juvenile age: Week 4

| **Fixed effect** | Estimate | *SE* | *F* | *df* | *df.res* | *P* |
|---|---|---|---|---|---|---|
| Intercept (freshwater) | 14.605 | 0.143 | 10381.640 | 1 | 142.970 | **<0.001** |
| Environment (fluctuating) | -0.512 | 0.119 | 251.920 | 2 | 760.040 | **<0.001** |
| Environment (stable) | 1.578 | 0.119 | | | | |

| **Random effect** | Variance | *sd* | Number of groups |
|---|---|---|---|
| Brood ID | 1.259 | 1.122 | 91 |
| Residual | 1.262 | 1.123 | |

| **Pairwise comparison** | *P* |
|---|---|
| Freshwater-Fluctuating | **<0.001** |
| Freshwater-Stable | **<0.001** |
| Stable-Fluctuating | **<0.001** |

(v) Juvenile age: Week 6

| **Fixed effect** | Estimate | *SE* | *F* | *df* | *df.res* | *P* |
|---|---|---|---|---|---|---|
| Intercept (freshwater) | 17.459 | 0.156 | 12560.720 | 1 | 182.870 | **<0.001** |
| Environment (fluctuating) | -1.025 | 0.156 | 176.640 | 2 | 728.630 | **<0.001** |
| Environment (stable) | 1.371 | 0.158 | | | | |

| **Random effect** | Variance | *sd* | Number of groups |
|---|---|---|---|
| Brood ID | 1.118 | 1.057 | 91 |
| Residual | 2.138 | 1.462 | |

| **Pairwise comparison** | *P* |
|---|---|
| Freshwater-Fluctuating | **<0.001** |
| Freshwater-Stable | **<0.001** |
| Stable-Fluctuating | **<0.001** |

### (B) Growth of adult females

(i) Model with environment*age interactive effect

| **Fixed effect** | Estimate | *SE* | *F* | *df* | *df.res* | *P* |
|---|---|---|---|---|---|---|
| Intercept (freshwater) | 17.495 | 0.252 | 4806.835 | 1 | 704.160 | **<0.001** |
| Environment (stable) | 0.620 | 0.235 | 3.548 | 2 | 369.750 | **0.030** |
| Environment (fluctuating) | 0.266 | 0.240 | | | | |
| Age | -0.264 | 0.035 | 58.207 | 1 | 1481.260 | **<0.001** |
| $Age^2$ | 4.092 | 0.156 | 686.216 | 1 | 1483.570 | **<0.001** |
| Environment (stable) * Age | 0.033 | 0.015 | 7.911 | 2 | 1487.990 | **<0.001** |
| Environment (fluctuating) * Age | -0.027 | 0.015 | | | | |

| **Random effect** | Variance | *sd* | Number of groups |
|---|---|---|---|
| Female ID | 1.992 | 1.412 | 319 |
| Brood ID | 0.897 | 0.947 | 96 |
| Residual | 1.007 | 1.003 | |

Due to a significant environment*age interaction, environment effects were tested for separately at each age:

(ii) Female adult age: Week 1 after maturity

| **Fixed effect** | Estimate | *SE* | *F* | *df* | *df.res* | *P* |
|---|---|---|---|---|---|---|
| Intercept (freshwater) | 21.405 | 0.171 | 15560.796 | 1 | 199.800 | **<0.001** |
| Environment (stable) | 0.724 | 0.186 | 7.680 | 2 | 260.140 | **<0.001** |
| Environment (fluctuating) | 0.336 | 0.189 | | | | |

| **Random effect** | Variance | *sd* | Number of groups |
|---|---|---|---|
| Brood ID | 1.113 | 1.055 | 96 |
| Residual | 1.531 | 1.237 | |

| **Pairwise comparison** | *P* |
|---|---|
| Freshwater-Stable | **<0.001** |
| Freshwater-Fluctuating | 0.182 |
| Stable-Fluctuating | 0.069 |

(iii) Female adult age: Week 3 after maturity

| **Fixed effect** | Estimate | *SE* | *F* | *df* | *df.res* | *P* |
|---|---|---|---|---|---|---|
| Intercept (freshwater) | 23.698 | 0.193 | 15089.085 | 1 | 202.160 | **<0.001** |
| Environment (stable) | 0.668 | 0.216 | 4.963 | 2 | 252.220 | **0.008** |
| Environment (fluctuating) | 0.262 | 0.219 | | | | |

| **Random effect** | Variance | *sd* | Number of groups |
|---|---|---|---|
| Brood ID | 1.267 | 1.126 | 95 |
| Residual | 2.022 | 1.422 | |

| **Pairwise comparison** | *P* |
|---|---|
| Freshwater-Stable | **0.006** |
| Freshwater-Fluctuating | 0.457 |
| Stable-Fluctuating | 0.118 |

(iv) Female adult age: Week 5 after maturity

| **Fixed effect** | Estimate | *SE* | *F* | *df* | *df.res* | *P* |
|---|---|---|---|---|---|---|
| Intercept (freshwater) | 25.180 | 0.206 | 14915.214 | 1 | 219.720 | **<0.001** |
| Environment (stable) | 0.762 | 0.249 | 5.846 | 2 | 254.110 | **0.003** |
| Environment (fluctuating) | 0.092 | 0.254 | | | | |

| **Random effect** | Variance | *sd* | Number of groups |
|---|---|---|---|
| Brood ID | 0.990 | 0.995 | 95 |
| Residual | 2.755 | 1.660 | |

| **Pairwise comparison** | *P* |
|---|---|
| Freshwater-Stable | **0.007** |
| Freshwater-Fluctuating | 0.931 |
| Stable-Fluctuating | **0.016** |

(v) Female adult age: Week 7 after maturity

| **Fixed effect** | Estimate | *SE* | *F* | *df* | *df.res* | *P* |
|---|---|---|---|---|---|---|
| Intercept (freshwater) | 26.674 | 0.221 | 14485.765 | 1 | 211.270 | **<0.001** |
| Environment (stable) | 0.998 | 0.264 | 9.171 | 2 | 245.840 | **<0.001** |
| Environment (fluctuating) | 0.081 | 0.270 | | | | |

| **Random effect** | Variance | *sd* | Number of groups |
|---|---|---|---|
| Brood ID | 1.269 | 1.126 | 95 |
| Residual | 3.035 | 1.742 | |

| **Pairwise comparison** | *P* |
|---|---|
| Freshwater-Stable | **<0.001** |
| Freshwater-Fluctuating | 0.952 |
| Stable-Fluctuating | **0.001** |

(vi) Female adult age: Week 9 after maturity

| **Fixed effect** | Estimate | *SE* | *F* | *df* | *df.res* | *P* |
|---|---|---|---|---|---|---|
| Intercept (freshwater) | 27.464 | 0.230 | 14167.388 | 1 | 204.020 | **<0.001** |
| Environment (stable) | 0.966 | 0.273 | 10.111 | 2 | 241.230 | **<0.001** |
| Environment (fluctuating) | -0.128 | 0.277 | | | | |

| **Random effect** | Variance | *sd* | Number of groups |
|---|---|---|---|
| Brood ID | 1.510 | 1.229 | 95 |
| Residual | 3.132 | 1.770 | |

| **Pairwise comparison** | *P* |
|---|---|
| Freshwater-Stable | **0.001** |
| Freshwater-Fluctuating | 0.890 |
| Stable-Fluctuating | **<0.001** |

(vii) Female adult age: Week 13 after maturity

| **Fixed effect** | Estimate | *SE* | *F* | *df* | *df.res* | *P* |
|---|---|---|---|---|---|---|
| Intercept (freshwater) | 28.988 | 0.242 | 14250.316 | 1 | 201.090 | **<0.001** |
| Environment (stable) | 0.856 | 0.289 | 8.428 | 2 | 235.850 | **<0.001** |
| Environment (fluctuating) | -0.261 | 0.297 | | | | |

| **Random effect** | Variance | *sd* | Number of groups |
|---|---|---|---|
| Brood ID | 1.645 | 1.282 | 95 |
| Residual | 3.485 | 1.867 | |

| **Pairwise comparison** | *P* |
|---|---|
| Freshwater-Stable | **0.010** |
| Freshwater-Fluctuating | 0.656 |
| Stable-Fluctuating | **<0.001** |

### (C) Growth of adult males

(i) Model with environment*age interactive effect

| **Fixed effect** | Estimate | *SE* | *F* | *df* | *df.res* | *P* |
|---|---|---|---|---|---|---|
| Intercept (freshwater) | 23.340 | 0.171 | 18553.540 | 1 | 267.660 | **<0.001** |
| Environment (stable) | 0.279 | 0.204 | 80.463 | 2 | 291.320 | **<0.001** |
| Environment (fluctuating) | -1.952 | 0.200 | | | | |
| Age | 0.095 | 0.009 | 119.042 | 1 | 1472.490 | **<0.001** |
| $Age^2$ | -0.291 | 0.041 | 49.798 | 1 | 1472.860 | **<0.001** |
| Environment (stable) * Age | 0.011 | 0.004 | 5.869 | 2 | 1473.570 | **0.003** |
| Environment (fluctuating) * Age | 0.013 | 0.004 | | | | |

| **Random effect** | Variance | *sd* | Number of groups |
|---|---|---|---|
| Male ID | 1.820 | 1.349 | 327 |
| Brood ID | 0.727 | 0.853 | 98 |
| Residual | 0.097 | 0.311 | |

Due to a significant environment*age interaction, environment effects were tested for separately at each age:

(ii) Male adult age: Week 1 after maturity

| **Fixed effect** | Estimate | *SE* | *F* | *df* | *df.res* | *P* |
|---|---|---|---|---|---|---|
| Intercept (freshwater) | 23.293 | 0.179 | 16830.178 | 1 | 222.620 | **<0.001** |
| Environment (stable) | 0.261 | 0.217 | 76.007 | 2 | 279.750 | **<0.001** |
| Environment (fluctuating) | -2.030 | 0.213 | | | | |

| **Random effect** | Variance | *sd* | Number of groups |
|---|---|---|---|
| Brood ID | 0.947 | 0.973 | 98 |
| Residual | 2.094 | 1.447 | |

| **Pairwise comparison** | *P* |
|---|---|
| Freshwater-Stable | 0.455 |
| Freshwater-Fluctuating | **<0.001** |
| Stable-Fluctuating | **<0.001** |

(iii) Male adult age: Week 3 after maturity

| **Fixed effect** | Estimate | *SE* | *F* | *df* | *df.res* | *P* |
|---|---|---|---|---|---|---|
| Intercept (freshwater) | 22.906 | 0.168 | 18540.847 | 1 | 216.070 | **<0.001** |
| Environment (stable) | 0.335 | 0.206 | 76.859 | 2 | 262.370 | **<0.001** |
| Environment (fluctuating) | -1.918 | 0.203 | | | | |

| **Random effect** | Variance | *sd* | Number of groups |
|---|---|---|---|
| Brood ID | 0.806 | 0.898 | 97 |
| Residual | 1.841 | 1.357 | |

| **Pairwise comparison** | *P* |
|---|---|
| Freshwater-Stable | 0.240 |
| Freshwater-Fluctuating | **<0.001** |
| Stable-Fluctuating | **<0.001** |

(iv) Male adult age: Week 5 after maturity

| **Fixed effect** | Estimate | *SE* | *F* | *df* | *df.res* | *P* |
|---|---|---|---|---|---|---|
| Intercept (freshwater) | 23.088 | 0.171 | 18178.328 | 1 | 223.260 | **<0.001** |
| Environment (stable) | 0.429 | 0.215 | 73.453 | 2 | 260.800 | **<0.001** |
| Environment (fluctuating) | -1.860 | 0.208 | | | | |

| **Random effect** | Variance | *sd* | Number of groups |
|---|---|---|---|
| Brood ID | 0.717 | 0.847 | 96 |
| Residual | 1.939 | 1.392 | |

| **Pairwise comparison** | *P* |
|---|---|
| Freshwater-Stable | 0.118 |
| Freshwater-Fluctuating | **<0.001** |
| Stable-Fluctuating | **<0.001** |

(v) Male adult age: Week 7 after maturity

| Fixed effect | Estimate | *SE* | *F* | *df* | *df.res* | *P* |
|---|---|---|---|---|---|---|
| Intercept (freshwater) | 23.220 | 0.174 | 17746.717 | 1 | 220.890 | **<0.001** |
| Environment (stable) | 0.455 | 0.220 | 70.188 | 2 | 253.260 | **<0.001** |
| Environment (fluctuating) | -1.831 | 0.213 | | | | |

| Random effect | Variance | *sd* | Number of groups |
|---|---|---|---|
| Brood ID | 0.687 | 0.829 | 95 |
| Residual | 1.935 | 1.391 | |

| Pairwise comparison | *P* |
|---|---|
| Freshwater-Stable | 0.101 |
| Freshwater-Fluctuating | **<0.001** |
| Stable-Fluctuating | **<0.001** |

(vi) Male adult age: Week 9 after maturity

| Fixed effect | Estimate | *SE* | *F* | *df* | *df.res* | *P* |
|---|---|---|---|---|---|---|
| Intercept (freshwater) | 23.435 | 0.172 | 18583.097 | 1 | 218.630 | **<0.001** |
| Environment (stable) | 0.419 | 0.220 | 69.178 | 2 | 249.460 | **<0.001** |
| Environment (fluctuating) | -1.815 | 0.211 | | | | |

| Random effect | Variance | *sd* | Number of groups |
|---|---|---|---|
| Brood ID | 0.646 | 0.804 | 94 |
| Residual | 1.857 | 1.363 | |

| Pairwise comparison | *P* |
|---|---|
| Freshwater-Stable | 0.142 |
| Freshwater-Fluctuating | **<0.001** |
| Stable-Fluctuating | **<0.001** |

(vii) Male adult age: Week 15 after maturity

| **Fixed effect** | Estimate | *SE* | *F* | *df* | *df.res* | *P* |
|---|---|---|---|---|---|---|
| Intercept (freshwater) | 23.566 | 0.173 | 18430.711 | 1 | 217.890 | **<0.001** |
| Environment (stable) | 0.423 | 0.224 | 66.557 | 2 | 238.520 | **<0.001** |
| Environment (fluctuating) | -1.803 | 0.214 | | | | |

| **Random effect** | Variance | *sd* | Number of groups |
|---|---|---|---|
| Brood ID | 0.618 | 0.786 | 94 |
| Residual | 1.854 | 1.362 | |

| **Pairwise comparison** | *P* |
|---|---|
| Freshwater-Stable | 0.146 |
| Freshwater-Fluctuating | **<0.001** |
| Stable-Fluctuating | **<0.001** |

**Table S2. Statistical outputs of environment effect on age at maturity of (A) females and (B) males.**

#### (A) Females

| **Fixed effect** | Estimate | *SE* | *F* | *df* | *df.res* | *P* |
|---|---|---|---|---|---|---|
| Intercept (freshwater) | 63.586 | 1.233 | 2655.730 | 1 | 228.420 | **<0.001** |
| Environment (stable) | -6.546 | 1.179 | 110.170 | 2 | 657.850 | **<0.001** |
| Environment (fluctuating) | 9.206 | 1.222 | | | | |

| **Random effect** | Variance | *sd* | Number of groups |
|---|---|---|---|
| Brood ID | 88.390 | 9.402 | 116 |
| Residual | 130.730 | 11.434 | |

| **Pairwise comparison** | *P* |
|---|---|
| Freshwater-Stable | **<0.001** |
| Freshwater-Fluctuating | **<0.001** |
| Stable-Fluctuating | **<0.001** |

#### (B) Males

| **Fixed effect** | Estimate | *SE* | *F* | *df* | *df.res* | *P* |
|---|---|---|---|---|---|---|
| Intercept (freshwater) | 89.429 | 1.537 | 3377.121 | 1 | 251.340 | **<0.001** |
| Environment (stable) | -7.020 | 1.576 | 18.962 | 2 | 601.220 | **<0.001** |
| Environment (fluctuating) | -9.551 | 1.567 | | | | |

| **Random effect** | Variance | *sd* | Number of groups |
|---|---|---|---|
| Brood ID | 117.100 | 10.820 | 116 |
| Residual | 206.500 | 14.370 | |

| **Pairwise comparison** | *P* |
|---|---|
| Freshwater-Stable | **<0.001** |
| Freshwater-Fluctuating | **<0.001** |
| Stable-Fluctuating | 0.168 |

**Table S3. Statistical outputs of environment effect on size at maturity of (A) females and (B) males.**

**(A) Females**

| **Fixed effect** | Estimate | *SE* | *F* | *df* | *df.res* | *P* |
|---|---|---|---|---|---|---|
| Intercept (freshwater) | 21.266 | 0.145 | 21625.717 | 1 | 234.750 | **<0.001** |
| Environment (stable) | 0.428 | 0.141 | 4.704 | 2 | 655.640 | **0.009** |
| Environment (fluctuating) | 0.210 | 0.146 | | | | |

| **Random effect** | Variance | *sd* | Number of groups |
|---|---|---|---|
| Brood ID | 1.156 | 1.075 | 116 |
| Residual | 1.874 | 1.369 | |

| **Pairwise comparison** | *P* |
|---|---|
| Freshwater-Stable | **0.007** |
| Freshwater-Fluctuating | 0.326 |
| Stable-Fluctuating | 0.202 |

**(B) Males**

| **Fixed effect** | Estimate | *SE* | *F* | *df* | *df.res* | *P* |
|---|---|---|---|---|---|---|
| Intercept (freshwater) | 23.180 | 0.147 | 24744 | 1 | 256.930 | **<0.001** |
| Environment (stable) | 0.281 | 0.155 | 174.670 | 2 | 589.110 | **<0.001** |
| Environment (fluctuating) | -2.144 | 0.154 | | | | |

| **Random effect** | Variance | *sd* | Number of groups |
|---|---|---|---|
| Brood ID | 0.997 | 0.999 | 116 |
| Residual | 2.006 | 1.417 | |

| **Pairwise comparison** | *P* |
|---|---|
| Freshwater-Stable | 0.168 |
| Freshwater-Fluctuating | **<0.001** |
| Stable-Fluctuating | **<0.001** |

**Table S4. Statistical outputs for the effects of developmental environment and adult age on female life-history and reproductive traits**

**(A) Adult mortality of females**

| Fixed effect | Estimated coefficient | *SE* | Hazard ratio | $\chi^2$ | *df* | *P* |
|---|---|---|---|---|---|---|
| Environment (stable) | 1.249 | 0.569 | 3.485 | 5.110 | 2 | 0.078 |
| Environment (fluctuating) | 1.182 | 0.578 | 3.262 | | | |

| Random effect | Variance | *sd* |
|---|---|---|
| Brood ID (intercept) | 0.492 | 0.702 |

**(B) Relative telomere length**

(i) Model with environment*age interactive effect

| **Fixed effect** | Estimate | *SE* | *F* | *df* | *df.res* | *P* |
|---|---|---|---|---|---|---|
| Intercept (freshwater, old) | 0.601 | 0.055 | 116.244 | 1 | 46.619 | **<0.001** |
| Environment (stable) | -0.100 | 0.045 | 2.725 | 2 | 116.000 | 0.070 |
| Environment (fluctuating) | -0.075 | 0.045 | | | | |
| Age (young) | -0.113 | 0.064 | 3.134 | 1 | 62.664 | 0.082 |
| Environment (**freshwater**) * Age (young) | 0.175 | 0.063 | 3.998 | 2 | 116.000 | **0.021** |
| Environment (fluctuating) * Age (young) | 0.121 | 0.063 | | | | |

| **Random effect** | Variance | *sd* | correlation | Number of groups |
|---|---|---|---|---|
| Brood ID (intercept) | 0.062 | 0.250 | | |
| Brood ID (young) | 0.060 | 0.245 | -0.950 | 35 |
| Residual | 0.030 | 0.174 | | |

(ii) Effect of absolute age (birth to age at sampling) on relative telomere length of females

| **Fixed effect** | Estimate | *SE* | *F* | *df* | *df.res* | *P* |
|---|---|---|---|---|---|---|
| Intercept (freshwater, old) | 0.702 | 0.091 | 56.788 | 1 | 144.100 | **<0.001** |
| Absolute age (standardized) | -0.104 | 0.074 | 1.835 | 1 | 136.360 | 0.178 |
| Environment (stable) | -0.115 | 0.046 | 3.201 | 2 | 120.690 | **0.044** |
| Environment (fluctuating) | -0.064 | 0.045 | | | | |
| Adult age (young) | -0.322 | 0.163 | 3.694 | 1 | 156.720 | 0.056 |
| Environment (stable) * Adult Age (young) | 0.187 | 0.064 | 4.547 | 2 | 116.040 | **0.013** |
| Environment (fluctuating) * Adult Age (young) | 0.133 | 0.064 | | | | |

| **Random effect** | Variance | *sd* | correlation | Number of groups |
|---|---|---|---|---|
| Brood ID (intercept) | 0.061 | 0.247 | | 35 |
| Brood ID (young) | 0.062 | 0.249 | -0.940 | |
| Residual | 0.030 | 0.173 | | |

Due to a significant environment*age interaction, environment effects were tested for separately at each age:

(iii) Relative telomere length of young females

| **Fixed effect** | Estimate | *SE* | *F* | *df* | *df.res* | *P* |
|---|---|---|---|---|---|---|
| Intercept (freshwater) | 0.489 | 0.034 | 208.246 | 1 | 78.535 | **<0.001** |
| Environment (stable) | 0.075 | 0.042 | 1.602 | 2 | 58.000 | 0.210 |
| Environment (fluctuating) | 0.045 | 0.042 | | | | |

| **Random effect** | Variance | *sd* | Number of groups |
|---|---|---|---|
| Brood ID | 0.008 | 0.089 | 30 |
| Residual | 0.026 | 0.163 | |

(iv) Relative telomere length of old females

| **Fixed effect** | Estimate | *SE* | *F* | *df* | *df.res* | *P* |
|---|---|---|---|---|---|---|
| Intercept (freshwater) | 0.599 | 0.056 | 113.429 | 1 | 47.516 | **<0.001** |
| Environment (stable) | -0.100 | 0.047 | 2.429 | 2 | 58.000 | 0.097 |
| Environment (fluctuating) | -0.075 | 0.047 | | | | |

| **Random effect** | Variance | *sd* | Number of groups |
|---|---|---|---|
| Brood ID | 0.061 | 0.248 | 30 |
| Residual | 0.034 | 0.184 | |

### (C) Relative gut length (log-transformed)

(i) Initial model with environment*age interactive effect

| **Fixed effect** | Estimate | *SE* | *F* | *df* | *df.res* | *P* |
|---|---|---|---|---|---|---|
| Intercept (freshwater, old) | 2.572 | 0.020 | 15883.71 | 1 | 352.12 | **<0.001** |
| Environment (stable) | -0.040 | 0.021 | 3.364 | 2 | 472.40 | **0.035** |
| Environment (fluctuating) | 0.014 | 0.022 | | | | |
| Age (young) | 0.052 | 0.034 | 2.296 | 1 | 455.09 | 0.130 |
| Log-transformed absolute body size (standardized) | 0.162 | 0.014 | 127.627 | 1 | 527.13 | **<0.001** |
| Environment (stable) * Age (young) | -0.055 | 0.031 | 1.754 | 2 | 507.27 | 0.174 |
| Environment (fluctuating) * Age (young) | -0.049 | 0.032 | | | | |

| **Random effect** | Variance | *sd* | correlation | Number of groups |
|---|---|---|---|---|
| Brood ID (intercept) | 0.003 | 0.056 | | 115 |
| Brood ID (young) | 0.007 | 0.085 | -0.940 | |
| Residual | 0.020 | 0.143 | | |

(ii) Final model excluding environment*age interaction to interpret the main effects

| **Fixed effect** | Estimate | *SE* | *F* | *df* | *df.res* | *P* |
|---|---|---|---|---|---|---|
| Intercept (freshwater, old) | 2.586 | 0.019 | 18486.128 | 1 | 334.720 | **<0.001** |
| Environment (stable) | -0.066 | 0.015 | 11.874 | 2 | 506.270 | **<0.001** |
| Environment (fluctuating) | -0.008 | 0.016 | | | | |
| Age (young) | 0.020 | 0.029 | 0.447 | 1 | 412.720 | 0.504 |
| Log-transformed absolute body size (standardized) | 0.164 | 0.014 | 131.942 | 1 | 526.990 | **<0.001** |

| **Random effect** | Variance | *sd* | correlation | Number of groups |
|---|---|---|---|---|
| Brood ID (intercept) | 0.003 | 0.057 | | 115 |
| Brood ID (young) | 0.008 | 0.088 | -0.950 | |
| Residual | 0.020 | 0.143 | | |

| **Pairwise comparison** | *P* |
|---|---|
| Freshwater-Stable | **<0.001** |
| Freshwater-Fluctuating | 0.855 |
| Stable-Fluctuating | **<0.001** |

(iii) Exclusion of the non-significant interactions did not significantly reduce model fit in the final model

| | No. parameter | AIC | BIC | Log-likelihood | Deviance | $\chi^2_2$ | *P* |
|---|---|---|---|---|---|---|---|
| Initial model (i) | 11 | -535.75 | -487.95 | 278.87 | -557.75 | | |
| Final model (ii) | 9 | -536.17 | -497.06 | 277.08 | -554.17 | 3.577 | 0.167 |

### (D) Immune response

(i) Initial model with environment*age interactive effect

| **Fixed effect** | Estimate | *SE* | $\chi^2$ | *df* | *P* |
|---|---|---|---|---|---|
| Intercept (freshwater, old) | -1.950 | 0.579 | 11.338 | 1 | **0.001** |
| Environment (stable) | 0.189 | 0.995 | 0.385 | 2 | 0.825 |
| Environment (fluctuating) | -0.095 | 0.675 | | | |
| Age (young) | -0.277 | 1.133 | 0.060 | 1 | 0.807 |
| Environment (stable) * Age (young) | -0.052 | 2.015 | 0.066 | 2 | 0.967 |
| Environment (fluctuating) * Age (young) | 0.108 | 1.073 | | | |

| **Random effect** | Variance | *sd* | Number of groups |
|---|---|---|---|
| Brood ID (intercept) | <0.001 | <0.001 | 115 |

(ii) Final model excluding environment*age interaction to interpret the main effects

| **Fixed effect** | Estimate | *SE* | $\chi^2$ | *df* | *P* |
|---|---|---|---|---|---|
| Intercept (freshwater, old) | -1.954 | 0.232 | 71.261 | 1 | **<0.001** |
| Environment (stable) | 0.164 | 0.284 | 0.639 | 2 | 0.726 |
| Environment (fluctuating) | -0.045 | 0.305 | | | |
| Age (young) | -0.265 | 0.237 | 1.253 | 1 | 0.263 |

| **Random effect** | Variance | *sd* | correlation | Number of groups |
|---|---|---|---|---|
| Brood ID (intercept) | <0.001 | <0.001 | | 115 |
| Brood ID (young) | <0.001 | 0.001 | -0.970 | |

Note: The two models are not comparable because of the difference in random factors (see main text)

### (E) Total egg number

(i) Initial model with environment*age interactive effect

| Fixed effect | Estimate | *SE* | $\chi^2$ | *df* | *P* |
|---|---|---|---|---|---|
| Intercept (freshwater, old) | 2.764 | 0.041 | 4498.274 | 1 | **<0.001** |
| Environment (stable) | 0.087 | 0.045 | 16.392 | 2 | **<0.001** |
| Environment (fluctuating) | -0.090 | 0.047 | | | |
| Age (young) | -0.937 | 0.069 | 186.933 | 1 | **<0.001** |
| Environment (stable) * Age (young) | -0.146 | 0.086 | 6.642 | 2 | **0.036** |
| Environment (fluctuating) * Age (young) | 0.048 | 0.091 | | | |

| Random effect | Variance | *sd* | correlation | Number of groups |
|---|---|---|---|---|
| Brood ID (intercept) | 0.062 | 0.249 | | 115 |
| Brood ID (young) | 0.043 | 0.207 | -0.700 | |

Due to a significant environment*age interaction, environment effects were tested for separately at each age:

(ii) Total egg number of young females

| Fixed effect | Estimate | *SE* | $\chi^2$ | *df* | *P* |
|---|---|---|---|---|---|
| Intercept (freshwater) | 1.800 | 0.056 | 1046.169 | 1 | **<0.001** |
| Environment (stable) | -0.025 | 0.068 | 0.193 | 2 | 0.908 |
| Environment (fluctuating) | -0.003 | 0.072 | | | |

| Random effect | Variance | *sd* | Number of groups |
|---|---|---|---|
| Brood ID | 0.048 | 0.218 | 110 |

(iii) Total egg number of old females

| **Fixed effect** | Estimate | *SE* | *F* | *df* | *df.res* | *P* |
|---|---|---|---|---|---|---|
| Intercept (freshwater) | 16.496 | 0.690 | 569.505 | 1 | 190.780 | **<0.001** |
| Environment (stable) | 1.701 | 0.783 | 7.517 | 2 | 229.780 | **<0.001** |
| Environment (fluctuating) | -1.261 | 0.804 | | | | |

| **Random effect** | Variance | *sd* | Number of groups |
|---|---|---|---|
| Brood ID | 16.470 | 4.058 | 95 |
| Residual | 24.910 | 4.991 | |

| **Pairwise comparison** | *P* |
|---|---|
| Freshwater-Stable | 0.079 |
| Freshwater-Fluctuating | 0.263 |
| Stable-Fluctuating | **<0.001** |

### (F) Egg size of young females

| **Fixed effect** | Estimate | *SE* | *F* | *df* | *df.res* | *P* |
|---|---|---|---|---|---|---|
| Intercept (freshwater) | 1.918 | 0.015 | 15372.239 | 1 | 281.1 | **<0.001** |
| Environment (stable) | 0.010 | 0.019 | 12.642 | 2 | 258.7 | **<0.001** |
| Environment (fluctuating) | -0.074 | 0.020 | | | | |

| **Random effect** | Variance | *sd* | Number of groups |
|---|---|---|---|
| Brood ID | 0.001 | 0.035 | 110 |
| Residual | 0.018 | 0.134 | |

| **Pairwise comparison** | *P* |
|---|---|
| Freshwater-Stable | 0.855 |
| Freshwater-Fluctuating | **0.001** |
| Stable-Fluctuating | **<0.001** |

### (G) Embryo number of old females

(i) Zero-inflation part of the model

| **Fixed effect** | Estimate | *SE* | $\chi^2$ | *df* | *P* |
|---|---|---|---|---|---|
| Intercept (freshwater) | -0.700 | 0.251 | 7.757 | 1 | **0.005** |
| Environment (stable) | -0.907 | 0.366 | 6.145 | 2 | **0.046** |
| Environment (fluctuating) | -0.345 | 0.347 | | | |

| **Random effect** | Variance | *sd* | Number of groups |
|---|---|---|---|
| Brood ID | 0.546 | 0.739 | 95 |

| **Pairwise comparison** | *P* |
|---|---|
| Freshwater-Stable | **0.037** |
| Freshwater-Fluctuating | 0.580 |
| Stable-Fluctuating | 0.290 |

(ii) Conditional part of the model

| **Fixed effect** | Estimate | *SE* | $\chi^2$ | *df* | *P* |
|---|---|---|---|---|---|
| Intercept (freshwater) | 2.693 | 0.062 | 1894.261 | 1 | **<0.001** |
| Environment (stable) | 0.137 | 0.075 | 6.755 | 2 | **0.034** |
| Environment (fluctuating) | -0.042 | 0.081 | | | |

| **Random effect** | Variance | *sd* | Number of groups |
|---|---|---|---|
| Brood ID | 0.065 | 0.256 | 95 |

| **Pairwise comparison** | *P* |
|---|---|
| Freshwater-Stable | 0.161 |
| Freshwater-Fluctuating | 0.862 |
| Stable-Fluctuating | **0.042** |

### (H) Likelihood of giving birth

| **Fixed effect** | Estimate | *SE* | $\chi^2$ | *df* | *P* |
|---|---|---|---|---|---|
| Intercept (freshwater) | -1.881 | 0.331 | 32.291 | 1 | **<0.001** |
| Environment (stable) | -0.116 | 0.432 | 0.759 | 2 | 0.684 |
| Environment (fluctuating) | -0.407 | 0.473 | | | |

| **Random effect** | Variance | *sd* | Number of groups |
|---|---|---|---|
| Brood ID | 0.326 | 0.571 | 95 |

### (I) Total offspring number

#### (i) Zero-inflation part of the model

| **Fixed effect** | Estimate | *SE* | $\chi^2$ | *df* | *P* |
|---|---|---|---|---|---|
| Intercept (freshwater) | -0.898 | 0.248 | 13.117 | 1 | **<0.001** |
| Environment (stable) | -0.688 | 0.355 | 3.867 | 2 | 0.145 |
| Environment (fluctuating) | -0.189 | 0.339 | | | |

| **Random effect** | Variance | *sd* | Number of groups |
|---|---|---|---|
| Brood ID | 0.212 | 0.461 | 95 |

#### (ii) Conditional part of the model

| **Fixed effect** | Estimate | *SE* | $\chi^2$ | *df* | *P* |
|---|---|---|---|---|---|
| Intercept (freshwater) | 2.684 | 0.067 | 1626.094 | 1 | **<0.001** |
| Environment (stable) | 0.198 | 0.083 | 7.865 | 2 | **0.020** |
| Environment (fluctuating) | -0.001 | 0.090 | | | |

| **Random effect** | Variance | *sd* | Number of groups |
|---|---|---|---|
| Brood ID | 0.058 | 0.240 | 95 |

| **Pairwise comparison** | *P* |
|---|---|
| Freshwater-Stable | **0.047** |
| Freshwater-Fluctuating | 0.999 |
| Stable-Fluctuating | 0.051 |

**Table S5. Statistical outputs for the effects of developmental environment and adult age on male life-history and reproductive traits**

**(A) Adult mortality of males**

| Fixed effect | Estimated coefficient | *SE* | Hazard ratio | $\chi^2$ | *df* | *P* |
|---|---|---|---|---|---|---|
| Environment (stable) | 0.068 | 0.343 | 1.070 | 0.477 | 2 | 0.788 |
| Environment (fluctuating) | -0.167 | 0.348 | 0.846 | | | |

| Random effect | Variance | *sd* |
|---|---|---|
| Brood ID (intercept) | <0.001 | 0.020 |

**(B) Relative telomere length**

(i) Initial model with environment*age interactive effect

| **Fixed effect** | Estimate | *SE* | *F* | *df* | *df.res* | *P* |
|---|---|---|---|---|---|---|
| Intercept (freshwater, old) | 0.730 | 0.046 | 251.463 | 1 | 64.831 | **<0.001** |
| Environment (stable) | -0.086 | 0.046 | 1.998 | 2 | 116.000 | 0.140 |
| Environment (fluctuating) | -0.072 | 0.046 | | | | |
| Age (young) | 0.079 | 0.054 | 2.047 | 1 | 89.701 | 0.156 |
| Environment (stable) * Age (young) | -0.087 | 0.065 | 1.140 | 2 | 116.000 | 0.323 |
| Environment (fluctuating) * Age (young) | -0.004 | 0.065 | | | | |

| **Random effect** | Variance | *sd* | correlation | Number of groups |
|---|---|---|---|---|
| Brood ID (intercept) | 0.032 | 0.180 | | 43 |
| Brood ID (young) | 0.019 | 0.137 | -0.870 | |
| Residual | 0.032 | 0.178 | | |

(ii) Final model excluding the non-significant interactive effect to interpret the main effects

| **Fixed effect** | Estimate | *SE* | *F* | *df* | *df.res* | *P* |
|---|---|---|---|---|---|---|
| Intercept (freshwater, old) | 0.745 | 0.042 | 314.073 | 1 | 47.051 | **<0.001** |
| Environment (stable) | -0.129 | 0.033 | 7.908 | 2 | 118.000 | **<0.001** |
| Environment (fluctuating) | -0.074 | 0.033 | | | | |
| Adult age (young) | 0.049 | 0.039 | 1.451 | 1 | 29.927 | 0.238 |

| **Random effect** | Variance | *sd* | correlation | Number of groups |
|---|---|---|---|---|
| Brood ID (intercept) | 0.032 | 0.180 | | 43 |
| Brood ID (young) | 0.019 | 0.137 | -0.870 | |
| Residual | 0.032 | 0.178 | | |

| **Pairwise comparison** | *P* |
|---|---|
| Freshwater-Stable | **<0.001** |
| Freshwater-Fluctuating | 0.065 |
| Stable-Fluctuating | 0.210 |

(iii) Exclusion of the non-significant interactions did not significantly reduce model fit in the final model

| | No. parameter | AIC | BIC | Log-likelihood | Deviance | $\chi^2_2$ | *P* |
|---|---|---|---|---|---|---|---|
| Initial model (i) | 10 | -39.8 | -7.9 | 29.9 | -59.8 | 2.336 | 0.311 |
| Final model (ii) | 8 | -41.5 | -15.9 | 28.7 | -57.5 | | |

(iv) Effect of absolute age (birth to age at sampling) on relative telomere length of females

| **Fixed effect** | Estimate | *SE* | *F* | *df* | *df.res* | *P* |
|---|---|---|---|---|---|---|
| Intercept (freshwater, old) | 0.768 | 0.071 | 110.560 | 1 | 146.070 | **<0.001** |
| Absolute age (standardized) | -0.021 | 0.055 | 0.136 | 1 | 161.980 | 0.713 |
| Environment (stable) | -0.132 | 0.034 | 7.637 | 2 | 121.360 | **<0.001** |
| Environment (fluctuating) | -0.078 | 0.035 | | | | |
| Adult age (young) | 0.009 | 0.111 | 0.006 | 1 | 170.580 | 0.938 |

| **Random effect** | Variance | *sd* | correlation | Number of groups |
|---|---|---|---|---|
| Brood ID (intercept) | 0.031 | 0.177 | | 43 |
| Brood ID (young) | 0.018 | 0.136 | -0.870 | |
| Residual | 0.032 | 0.179 | | |

#### (C) Relative gut length (log-transformed)

(i) Initial model with environment*age interactive effect

| **Fixed effect** | Estimate | *SE* | *F* | *df* | *df.res* | *P* |
|---|---|---|---|---|---|---|
| Intercept (freshwater, old) | 2.188 | 0.014 | 25305.46 | 1 | 275.31 | **<0.001** |
| Environment (stable) | -0.003 | 0.018 | 0.029 | 2 | 435.23 | 0.971 |
| Environment (fluctuating) | 0.002 | 0.019 | | | | |
| Age (young) | 0.082 | 0.019 | 17.923 | 1 | 323.07 | **<0.001** |
| Log-transformed absolute body size (standardized) | 0.075 | 0.007 | 124.718 | 1 | 411.49 | **<0.001** |
| Environment (stable) * Age (young) | 0.014 | 0.027 | 0.553 | 2 | 453.97 | 0.576 |
| Environment (fluctuating) * Age (young) | 0.027 | 0.026 | | | | |

| **Random effect** | Variance | *sd* | correlation | Number of groups |
|---|---|---|---|---|
| Brood ID (intercept) | 0.001 | 0.038 | | 114 |
| Brood ID (young) | 0.001 | 0.030 | -0.95 | |
| Residual | 0.014 | 0.117 | | |

(ii) Final model excluding the non-significant interactive effect to interpret the main effects

| **Fixed effect** | Estimate | *SE* | *F* | *df* | *df.res* | *P* |
|---|---|---|---|---|---|---|
| Intercept (freshwater, old) | 2.182 | 0.012 | 33992.840 | 1 | 240.370 | **<0.001** |
| Environment (stable) | 0.004 | 0.013 | 0.482 | 2 | 472.650 | 0.618 |
| Environment (fluctuating) | 0.014 | 0.015 | | | | |
| Adult age (young) | 0.097 | 0.011 | 72.657 | 1 | 96.610 | **<0.001** |
| Log-transformed absolute body size (standardized) | 0.075 | 0.007 | 124.002 | 1 | 409.44 | **<0.001** |

| **Random effect** | Variance | *sd* | correlation | Number of groups |
|---|---|---|---|---|
| Brood ID (intercept) | 0.001 | 0.037 | | 114 |
| Brood ID (young) | 0.001 | 0.031 | -0.92 | |
| Residual | 0.014 | 0.117 | | |

(iii) Exclusion of the non-significant interactions did not significantly reduce model fit in the final model

| | No. parameter | AIC | BIC | Log-likelihood | Deviance | $\chi^2_2$ | *P* |
|---|---|---|---|---|---|---|---|
| Initial model (i) | 11 | -705.10 | -658.37 | 363.55 | -727.10 | 1.126 | 0.570 |
| Final model (ii) | 9 | -707.98 | -669.74 | 362.99 | -725.98 | | |

**(D) Immune response**

(i) Initial model with environment*age interactive effect

| **Fixed effect** | Estimate | *SE* | $\chi^2$ | *df* | *P* |
|---|---|---|---|---|---|
| Intercept (freshwater, old) | -1.799 | 0.259 | 48.198 | 1 | **<0.001** |
| Environment (stable) | 0.296 | 0.347 | 3.656 | 2 | 0.161 |
| Environment (fluctuating) | -0.429 | 0.398 | | | |
| Age (young) | -0.138 | 0.413 | 0.111 | 1 | 0.739 |
| Environment (stable) * Age (young) | -0.228 | 0.554 | 0.349 | 2 | 0.840 |
| Environment (fluctuating) * Age (young) | 0.095 | 0.609 | | | |

| **Random effect** | Variance | *sd* | Number of groups |
|---|---|---|---|
| Brood ID (intercept) | <0.001 | <0.001 | 113 |

(ii) Final model excluding environment*age interaction to interpret the main effects

| **Fixed effect** | Estimate | *SE* | $\chi^2$ | *df* | *P* |
|---|---|---|---|---|---|
| Intercept (freshwater, old) | -1.773 | 0.220 | 64.760 | 1 | **<0.001** |
| Environment (stable) | 0.204 | 0.271 | 4.237 | 2 | 0.120 |
| Environment (fluctuating) | -0.384 | 0.301 | | | |
| Age (young) | -0.205 | 0.234 | 0.768 | 1 | 0.381 |

| **Random effect** | Variance | *sd* | correlation | Number of groups |
|---|---|---|---|---|
| Brood ID (intercept) | <0.001 | 0.001 | | 113 |
| Brood ID (young) | <0.001 | 0.001 | -0.930 | |

Note: The two models are not comparable because of the difference in random factors (see main text)

### (E) Number of mating attempts

(i) Zero-inflation part of the model with environment*age interactive effect

| **Fixed effect** | Estimate | *SE* | $\chi^2$ | *df* | *P* |
|---|---|---|---|---|---|
| Intercept (freshwater, old) | -14.460 | 3.941 | 13.462 | 1 | **<0.001** |
| Environment (stable) | 1.968 | 4.207 | 0.236 | 2 | 0.889 |
| Environment (fluctuating) | 1.102 | 4.617 | | | |
| Age (young) | 0.312 | 2.680 | 0.014 | 1 | 0.907 |
| Environment (stable) * Age (young) | -12.747 | 4.536 | 9.079 | 2 | **0.011** |
| Environment (fluctuating) * Age (young) | -13.845 | 6.601 | | | |

| **Random effect** | Variance | *sd* | Number of groups |
|---|---|---|---|
| Brood ID (intercept) | <0.001 | <0.001 | 98 |
| Male ID (intercept) | 1733.000 | 41.630 | 323 |

(ii) Conditional part of the model with environment*age interactive effect

| **Fixed effect** | Estimate | *SE* | $\chi^2$ | *df* | *P* |
|---|---|---|---|---|---|
| Intercept (freshwater, old) | 3.097 | 0.082 | 1440.625 | 1 | **<0.001** |
| Environment (stable) | 0.330 | 0.113 | 12.932 | 2 | **0.002** |
| Environment (fluctuating) | 0.353 | 0.107 | | | |
| Age (young) | 0.698 | 0.113 | 38.366 | 1 | **<0.001** |
| Environment (stable) * Age (young) | -0.423 | 0.157 | 8.498 | 2 | **0.014** |
| Environment (fluctuating) * Age (young) | -0.359 | 0.150 | | | |

| **Random effect** | Variance | *sd* | Number of groups |
|---|---|---|---|
| Brood ID (intercept) | 0.032 | 0.180 | 98 |
| Male ID (intercept) | <0.001 | 0.001 | 323 |

Due to a significant environment*age interaction, environment effects were tested for separately at each age:

(iii) Zero-inflation part of the model testing for the environment effect (young male)

| **Fixed effect** | Estimate | *SE* | $\chi^2$ | *df* | *P* |
|---|---|---|---|---|---|
| Intercept (freshwater) | -3.255 | 0.610 | 28.458 | 1 | **<0.001** |
| Environment (stable) | 0.073 | 0.811 | 1.619 | 2 | 0.445 |
| Environment (fluctuating) | -1.652 | 1.411 | | | |

| **Random effect** | Variance | *sd* | Number of groups |
|---|---|---|---|
| Brood ID (intercept) | <0.001 | <0.001 | 98 |

(iv) Conditional part of the model testing for the environment effect (young male)

| **Fixed effect** | Estimate | *SE* | $\chi^2$ | *df* | *P* |
|---|---|---|---|---|---|
| Intercept (freshwater) | 3.814 | 0.088 | 1885.711 | 1 | **<0.001** |
| Environment (stable) | -0.100 | 0.113 | 0.986 | 2 | 0.611 |
| Environment (fluctuating) | -0.014 | 0.109 | | | |

| **Random effect** | Variance | *sd* | Number of groups |
|---|---|---|---|
| Brood ID (intercept) | 0.015 | 0.124 | 98 |

(v) Zero-inflation part of the model testing for the environment effect (old male)

| **Fixed effect** | Estimate | *SE* | $\chi^2$ | *df* | *P* |
|---|---|---|---|---|---|
| Intercept (freshwater) | -5.652 | 3.282 | 2.965 | 1 | 0.085 |
| Environment (stable) | 2.550 | 3.322 | 0.878 | 2 | 0.645 |
| Environment (fluctuating) | 2.003 | 3.340 | | | |

(vi) Conditional part of the model testing for the environment effect (old male)

| **Fixed effect** | Estimate | *SE* | $\chi^2$ | *df* | *P* |
|---|---|---|---|---|---|
| Intercept (freshwater) | 3.117 | 0.086 | 1316.515 | 1 | **<0.001** |
| Environment (stable) | 0.330 | 0.117 | 11.367 | 2 | **0.003** |
| Environment (fluctuating) | 0.341 | 0.112 | | | |

| **Random effect** | Variance | *sd* | Number of groups |
|---|---|---|---|
| Brood ID (intercept) | 0.014 | 0.118 | 94 |

| **Pairwise comparison** | *P* |
|---|---|
| Freshwater-Stable | **0.015** |
| Freshwater-Fluctuating | **0.007** |
| Stable-Fluctuating | 0.994 |

#### (F) Likelihood of successful mating

(i) Initial model with environment*age interactive effect

| Fixed effect | Estimate | *SE* | $\chi^2$ | *df* | *P* |
|---|---|---|---|---|---|
| Intercept (freshwater, old) | -24.563 | 4.743 | 26.819 | 1 | **<0.001** |
| Environment (stable) | -19.524 | 5.343 | 19.212 | 2 | **<0.001** |
| Environment (fluctuating) | -0.125 | 3.419 | | | |
| Age (young) | 0.025 | 2.970 | <0.001 | 1 | 0.993 |
| Environment (stable) * Age (young) | 22.715 | 4.072 | 39.260 | 2 | **<0.001** |
| Environment (fluctuating) * Age (young) | 2.600 | 3.595 | | | |

| Random effect | Variance | *sd* | correlation | Number of groups |
|---|---|---|---|---|
| Brood ID (intercept) | 1107.000 | 33.270 | | 98 |
| Brood ID (young) | 4426.000 | 66.530 | -1.000 | |
| Male ID (intercept) | 3240.000 | 56.920 | | 323 |

| Pairwise comparison (age effect at each treatment) | *P* |
|---|---|
| Freshwater: young vs old | 0.993 |
| Stable: young vs old | **<0.001** |
| Fluctuating: young vs old | 0.535 |

Due to a significant environment*age interaction, environment effects were tested for separately at each age:

(ii) Young males: Likelihood of successful mating

| Fixed effect | Estimate | *SE* | $\chi^2$ | *df* | *P* |
|---|---|---|---|---|---|
| Intercept (freshwater) | -1.133 | 0.264 | 18.452 | 1 | **<0.001** |
| Environment (stable) | -0.022 | 0.356 | 0.040 | 2 | 0.980 |
| Environment (fluctuating) | -0.067 | 0.349 | | | |

| Random effect | Variance | *sd* | Number of groups |
|---|---|---|---|
| Brood ID (intercept) | <0.001 | <0.001 | 98 |

(iii) Old males: Likelihood of successful mating

| **Fixed effect** | Estimate | *SE* | $\chi^2$ | *df* | *P* |
|---|---|---|---|---|---|
| Intercept (freshwater) | -2.009 | 0.439 | 20.903 | 1 | **<0.001** |
| Environment (stable) | -0.959 | 0.549 | 3.061 | 2 | 0.216 |
| Environment (fluctuating) | -0.374 | 0.462 | | | |

| **Random effect** | Variance | *sd* | Number of groups |
|---|---|---|---|
| Brood ID (intercept) | 1.095 | 1.047 | 94 |

### (G) Time spent with female

(i) Initial model with environment*age interactive effect

| **Fixed effect** | Estimate | *SE* | *F* | *df* | *df.res* | *P* |
|---|---|---|---|---|---|---|
| Intercept (freshwater, old) | 353.330 | 20.000 | 308.969 | 1 | 264.410 | **<0.001** |
| Environment (stable) | 32.310 | 26.950 | 1.393 | 2 | 461.260 | 0.250 |
| Environment (fluctuating) | 42.330 | 25.790 | | | | |
| Age (young) | -20.650 | 25.800 | 0.633 | 1 | 275.380 | 0.427 |
| Environment (stable) * Age (young) | -35.810 | 33.830 | 1.241 | 2 | 255.000 | 0.291 |
| Environment (fluctuating) * Age (young) | -51.370 | 32.640 | | | | |

| **Random effect** | Variance | *sd* | correlation | Number of groups |
|---|---|---|---|---|
| Brood ID (intercept) | 5276 | 72.640 | | 98 |
| Brood ID (young) | 7433 | 86.210 | -0.970 | |
| Male ID (intercept) | 6942 | 83.320 | | 323 |
| Residual | 21686 | 147.260 | | |

(ii) Final model excluding the non-significant interactive effect to interpret the main effects

| **Fixed effect** | Estimate | *SE* | *F* | *df* | *df.res* | *P* |
|---|---|---|---|---|---|---|
| Intercept (freshwater, old) | 368.540 | 17.210 | 453.844 | 1 | 218.619 | **<0.001** |
| Environment (stable) | 14.020 | 20.290 | 0.367 | 2 | 287.991 | 0.694 |
| Environment (fluctuating) | 15.910 | 19.620 | | | | |
| Adult age (young) | -51.390 | 15.730 | 10.559 | 1 | 84.508 | **0.002** |

| **Random effect** | Variance | *sd* | correlation | Number of groups |
|---|---|---|---|---|
| Brood ID (intercept) | 4812 | 69.370 | | 98 |
| Brood ID (young) | 7083 | 84.160 | -0.960 | |
| Male ID (intercept) | 7112 | 84.330 | | 323 |
| Residual | 21634 | 147.080 | | |

(iii) Exclusion of the non-significant interactions did not significantly reduce model fit in the final model

| | No. parameter | AIC | BIC | Log-likelihood | Deviance | $\chi^2_2$ | *P* |
|---|---|---|---|---|---|---|---|
| Initial model (i) | 11 | 7372.7 | 7420.3 | -3675.4 | 7350.7 | 2.463 | 0.292 |
| Final model (ii) | 9 | 7371.2 | 7410.1 | -3676.6 | 7353.2 | | |

**(H) Sperm velocity- VCL**

(i) Model with environment*age interactive effect

| **Fixed effect** | Estimate | *SE* | *F* | *df* | *df.res* | *P* |
|---|---|---|---|---|---|---|
| Intercept (freshwater, old) | 166.466 | 2.369 | 4882.279 | 1 | 265.200 | **<0.001** |
| Environment (stable) | 2.603 | 3.222 | 0.331 | 2 | 443.230 | 0.719 |
| Environment (fluctuating) | 1.700 | 3.084 | | | | |
| Age (young) | 12.198 | 3.301 | 13.507 | 1 | 281.290 | **<0.001** |
| Environment (stable) * Age (young) | -9.648 | 4.378 | 5.478 | 2 | 256.720 | **0.005** |
| Environment (fluctuating) * Age (young) | -13.996 | 4.235 | | | | |

| **Random effect** | Variance | *sd* | correlation | Number of groups |
|---|---|---|---|---|
| Brood ID (intercept) | 63.690 | 7.981 | | 97 |
| Brood ID (young) | 99.450 | 9.972 | -0.110 | |
| Male ID (intercept) | 35.450 | 5.954 | | 317 |
| Residual | 379.680 | 19.485 | | |

Due to a significant environment*age interaction, environment effects were tested for separately at each age:

(ii) Sperm velocity of young males

| **Fixed effect** | Estimate | *SE* | *F* | *df* | *df.res* | *P* |
|---|---|---|---|---|---|---|
| Intercept (freshwater) | 178.592 | 2.775 | 4126.773 | 1 | 251.530 | **<0.001** |
| Environment (stable) | -7.043 | 3.428 | 7.014 | 2 | 224.100 | **0.001** |
| Environment (fluctuating) | -12.614 | 3.361 | | | | |

| **Random effect** | Variance | *sd* | Number of groups |
|---|---|---|---|
| Brood ID (intercept) | 135.000 | 11.620 | 97 |
| Residual | 451.000 | 21.240 | |

| **Pairwise comparison** | *P* |
|---|---|
| Freshwater-Stable | 0.103 |
| Freshwater-Fluctuating | **<0.001** |
| Stable-Fluctuating | 0.173 |

(iii) Sperm velocity of old males

| Fixed effect | Estimate | *SE* | *F* | *df* | *df.res* | *P* |
|---|---|---|---|---|---|---|
| Intercept (freshwater) | 167.038 | 2.323 | 5134.543 | 1 | 225.820 | **<0.001** |
| Environment (stable) | 1.698 | 3.129 | 0.146 | 2 | 246.040 | 0.864 |
| Environment (fluctuating) | 0.808 | 2.994 | | | | |

| Random effect | Variance | *sd* | Number of groups |
|---|---|---|---|
| Brood ID (intercept) | 72.220 | 8.498 | 94 |
| Residual | 381.180 | 19.524 | |

### (I) Total sperm count (log-transformed)

(i) Initial model with environment*age interactive effect

| Fixed effect | Estimate | *SE* | *F* | *df* | *df.res* | *P* |
|---|---|---|---|---|---|---|
| Intercept (freshwater, old) | 15.386 | 0.066 | 53427.953 | 1 | 234.570 | **<0.001** |
| Environment (stable) | -0.077 | 0.083 | 8.715 | 2 | 418.540 | **<0.001** |
| Environment (fluctuating) | -0.312 | 0.080 | | | | |
| Age (young) | -0.266 | 0.092 | 8.294 | 1 | 237.140 | **0.004** |
| Environment (stable) * Age (young) | -0.141 | 0.114 | 1.014 | 2 | 236.030 | 0.365 |
| Environment (fluctuating) * Age (young) | -0.143 | 0.110 | | | | |

| Random effect | Variance | *sd* | correlation | Number of groups |
|---|---|---|---|---|
| Brood ID (intercept) | 0.109 | 0.331 | | 97 |
| Brood ID (young) | 0.205 | 0.453 | -0.910 | |
| Male ID (intercept) | 0.036 | 0.190 | | 312 |
| Residual | 0.217 | 0.465 | | |

(ii) Final model excluding the non-significant interactive effect to interpret the main effects

| **Fixed effect** | Estimate | *SE* | *F* | *df* | *df.res* | *P* |
|---|---|---|---|---|---|---|
| Intercept (freshwater, old) | 15.430 | 0.059 | 67599.516 | 1 | 181.690 | **<0.001** |
| Environment (stable) | -0.146 | 0.062 | 21.534 | 2 | 262.170 | **<0.001** |
| Environment (fluctuating) | -0.383 | 0.059 | | | | |
| Adult age (young) | -0.357 | 0.066 | 29.055 | 1 | 90.276 | **<0.001** |

| **Random effect** | Variance | *sd* | correlation | Number of groups |
|---|---|---|---|---|
| Brood ID (intercept) | 0.119 | 0.345 | | 97 |
| Brood ID (young) | 0.235 | 0.485 | -0.920 | |
| Male ID (intercept) | 0.041 | 0.203 | | 312 |
| Residual | 0.208 | 0.456 | | |

| **Pairwise comparison** | *P* |
|---|---|
| Freshwater-Stable | 0.051 |
| Freshwater-Fluctuating | **<0.001** |
| Stable-Fluctuating | **<0.001** |

(iii) Exclusion of the non-significant interactions did not significantly reduce model fit in the final model

| | No. parameter | AIC | BIC | Log-likelihood | Deviance | $\chi^2_2$ | *P* |
|---|---|---|---|---|---|---|---|
| Initial model (i) | 11 | 887.1 | 934.0 | -432.6 | 865.1 | 1.841 | 0.398 |
| Final model (ii) | 9 | 885.0 | 923.3 | -433.5 | 867.0 | | |

**Table S6. Statistical outputs for the effects of environment, age and *body size* on female life-history and reproductive traits.** Two additional analyses were run to test if body size had an effect on the measured traits: (1) final model with absolute body length (globally standardized across all environments) as a covariate and (2) final model with within-group body length (standardized within each environment/age combination) as a covariate. Table S4 reported the final model for each female trait. *Methods* in the main text describe the details of the analysis (see *Analysis 2*).

#### (A) Relative telomere length

Due to a significant environment*age interaction, body size effects were tested for separately at each age:

(i) Effect of absolute body length across environments (young female)

| **Fixed effect** | Estimate | *SE* | *F* | *df* | *df.res* | *P* |
|---|---|---|---|---|---|---|
| Intercept (freshwater) | 0.488 | 0.034 | 206.168 | 1 | 78.214 | **<0.001** |
| Environment (stable) | 0.074 | 0.042 | 1.555 | 2 | 57.861 | 0.220 |
| Environment (fluctuating) | 0.048 | 0.043 | | | | |
| Absolute body size (standardized) | 0.009 | 0.020 | 0.159 | 1 | 85.691 | 0.691 |

| **Random effect** | Variance | *sd* | Number of groups |
|---|---|---|---|
| Brood ID | 0.008 | 0.088 | 30 |
| Residual | 0.027 | 0.164 | |

(ii) Effect of body length within each environment (young female)

| **Fixed effect** | Estimate | *SE* | *F* | *df* | *df.res* | *P* |
|---|---|---|---|---|---|---|
| Intercept (freshwater) | 0.489 | 0.034 | 207.017 | 1 | 78.193 | **<0.001** |
| Environment (stable) | 0.075 | 0.042 | 1.573 | 2 | 57.092 | 0.216 |
| Environment (fluctuating) | 0.045 | 0.042 | | | | |
| Within-group body size (standardized) | 0.008 | 0.020 | 0.156 | 1 | 85.699 | 0.694 |

| **Random effect** | Variance | *sd* | Number of groups |
|---|---|---|---|
| Brood ID | 0.008 | 0.088 | 30 |
| Residual | 0.027 | 0.164 | |

(iii) Effect of absolute body length across environments (old female)

| **Fixed effect** | Estimate | *SE* | *F* | *df* | *df.res* | *P* |
|---|---|---|---|---|---|---|
| Intercept (freshwater) | 0.596 | 0.057 | 110.330 | 1 | 46.897 | **<0.001** |
| Environment (stable) | -0.087 | 0.049 | 2.005 | 2 | 57.916 | 0.144 |
| Environment (fluctuating) | -0.078 | 0.047 | | | | |
| Absolute body size (standardized) | -0.027 | 0.027 | 1.018 | 1 | 72.537 | 0.316 |

| **Random effect** | Variance | *sd* | Number of groups |
|---|---|---|---|
| Brood ID | 0.063 | 0.250 | 30 |
| Residual | 0.033 | 0.183 | |

(iv) Effect of body length within each environment (old female)

| **Fixed effect** | Estimate | *SE* | *F* | *df* | *df.res* | *P* |
|---|---|---|---|---|---|---|
| Intercept (freshwater) | 0.599 | 0.057 | 112.244 | 1 | 46.606 | **<0.001** |
| Environment (stable) | -0.100 | 0.047 | 2.458 | 2 | 57.019 | 0.095 |
| Environment (fluctuating) | -0.075 | 0.047 | | | | |
| Within-group body size (standardized) | -0.029 | 0.026 | 1.171 | 1 | 73.388 | 0.283 |

| **Random effect** | Variance | *sd* | Number of groups |
|---|---|---|---|
| Brood ID | 0.063 | 0.250 | 30 |
| Residual | 0.033 | 0.183 | |

### (B) Immune response

(i) Effect of absolute body length across environments

| **Fixed effect** | Estimate | *SE* | $\chi^2$ | *df* | *P* |
|---|---|---|---|---|---|
| Intercept (freshwater, old) | -2.291 | 0.345 | 44.175 | 1 | **<0.001** |
| Environment (stable) | 0.130 | 0.285 | 0.341 | 2 | 0.843 |
| Environment (fluctuating) | -0.020 | 0.305 | | | |
| Age (young) | 0.398 | 0.540 | 0.545 | 1 | 0.460 |
| Absolute body size (standardized) | 0.368 | 0.267 | 1.897 | 1 | 0.168 |

| **Random effect** | Variance | *sd* | correlation | Number of groups |
|---|---|---|---|---|
| Brood ID (intercept) | <0.001 | <0.001 | | 115 |
| Brood ID (young) | <0.001 | <0.001 | -0.89 | |

(ii) Effect of body length within each environment/age combination

| **Fixed effect** | Estimate | *SE* | $\chi^2$ | *df* | *P* |
|---|---|---|---|---|---|
| Intercept (freshwater, old) | -1.968 | 0.232 | 71.779 | 1 | **<0.001** |
| Environment (stable) | 0.164 | 0.284 | 0.639 | 2 | 0.727 |
| Environment (fluctuating) | -0.045 | 0.305 | | | |
| Age (young) | -0.266 | 0.237 | 1.259 | 1 | 0.262 |
| Within-group body size (standardized) | 0.166 | 0.120 | 1.899 | 1 | 0.168 |

| **Random effect** | Variance | *sd* | correlation | Number of groups |
|---|---|---|---|---|
| Brood ID (intercept) | <0.001 | <0.001 | | 115 |
| Brood ID (young) | <0.001 | <0.001 | -0.91 | |

#### (C) Total egg number

Due to a significant environment*age interaction, body size effects were tested for separately at each age:

(i) Effect of absolute body length across environments (young female)

| Fixed effect | Estimate | *SE* | $\chi^2$ | *df* | *P* |
|---|---|---|---|---|---|
| Intercept (freshwater) | 1.749 | 0.052 | 1115.123 | 1 | **<0.001** |
| Environment (stable) | -0.031 | 0.064 | 1.251 | 2 | 0.535 |
| Environment (fluctuating) | 0.034 | 0.068 | | | |
| Absolute body size (standardized) | 0.244 | 0.027 | 82.332 | 1 | **<0.001** |

| Random effect | Variance | *sd* | Number of groups |
|---|---|---|---|
| Brood ID | 0.030 | 0.173 | 110 |

(ii) Effect of body length within each environment (young female)

| Fixed effect | Estimate | *SE* | $\chi^2$ | *df* | *P* |
|---|---|---|---|---|---|
| Intercept (freshwater) | 1.791 | 0.052 | 1194.059 | 1 | **<0.001** |
| Environment (stable) | -0.060 | 0.064 | 0.933 | 2 | 0.627 |
| Environment (fluctuating) | -0.050 | 0.068 | | | |
| Within-group body size (standardized) | 0.240 | 0.027 | 80.422 | 1 | **<0.001** |

| Random effect | Variance | *sd* | Number of groups |
|---|---|---|---|
| Brood ID | 0.030 | 0.174 | 110 |

(iii) Effect of absolute body length across environments (old female)

| Fixed effect | Estimate | *SE* | *F* | *df* | *df.res* | *P* |
|---|---|---|---|---|---|---|
| Intercept (freshwater) | 16.690 | 0.548 | 923.214 | 1 | 194.290 | **<0.001** |
| Environment (stable) | 0.326 | 0.644 | 1.582 | 2 | 234.590 | 0.208 |
| Environment (fluctuating) | -0.789 | 0.652 | | | | |
| Absolute body size (standardized) | 3.709 | 0.298 | 153.394 | 1 | 279.560 | **<0.001** |

| Random effect | Variance | *sd* | Number of groups |
|---|---|---|---|
| Brood ID | 9.644 | 3.106 | 95 |
| Residual | 16.487 | 4.060 | |

(iv) Effect of body length within each environment (old female)

| **Fixed effect** | Estimate | *SE* | *F* | *df* | *df.res* | *P* |
|---|---|---|---|---|---|---|
| Intercept (freshwater) | 16.256 | 0.549 | 872.052 | 1 | 193.690 | **<0.001** |
| Environment (stable) | 1.746 | 0.635 | 9.961 | 2 | 231.150 | **<0.001** |
| Environment (fluctuating) | -1.000 | 0.652 | | | | |
| Within-group body size (standardized) | 3.640 | 0.294 | 151.988 | 1 | 279.580 | **<0.001** |

| **Random effect** | Variance | *sd* | Number of groups |
|---|---|---|---|
| Brood ID | 9.721 | 3.118 | 95 |
| Residual | 16.522 | 4.065 | |

| **Pairwise comparison** | *P* |
|---|---|
| Freshwater-Stable | **0.018** |
| Freshwater-Fluctuating | 0.279 |
| Stable-Fluctuating | **<0.001** |

### (D) Egg size of young females

(i) Effect of absolute body length across environments

| **Fixed effect** | Estimate | *SE* | *F* | *df* | *df.res* | *P* |
|---|---|---|---|---|---|---|
| Intercept (freshwater) | 1.914 | 0.015 | 15611.253 | 1 | 280.99 | **<0.001** |
| Environment (stable) | 0.012 | 0.019 | 11.106 | 2 | 258.59 | **<0.001** |
| Environment (fluctuating) | -0.067 | 0.020 | | | | |
| Absolute body size (standardized) | 0.023 | 0.008 | 8.129 | 1 | 266.70 | **0.005** |

| **Random effect** | Variance | *sd* | Number of groups |
|---|---|---|---|
| Brood ID | 0.001 | 0.033 | 110 |
| Residual | 0.016 | 0.125 | |

| **Pairwise comparison** | *P* |
|---|---|
| Freshwater-Stable | 0.816 |
| Freshwater-Fluctuating | **0.003** |
| Stable-Fluctuating | **<0.001** |

(ii) Effect of body length within each environment

| **Fixed effect** | Estimate | *SE* | *F* | *df* | *df.res* | *P* |
|---|---|---|---|---|---|---|
| Intercept (freshwater) | 1.919 | 0.015 | 15889.453 | 1 | 280.59 | **<0.001** |
| Environment (stable) | 0.008 | 0.019 | 12.937 | 2 | 260.78 | **<0.001** |
| Environment (fluctuating) | -0.075 | 0.020 | | | | |
| Within-group body size (standardized) | 0.025 | 0.008 | 9.755 | 1 | 268.06 | **0.002** |

| **Random effect** | Variance | *sd* | Number of groups |
|---|---|---|---|
| Brood ID | 0.001 | 0.032 | 110 |
| Residual | 0.016 | 0.125 | |

| **Pairwise comparison** | *P* |
|---|---|
| Freshwater-Stable | 0.906 |
| Freshwater-Fluctuating | **<0.001** |
| Stable-Fluctuating | **<0.001** |

### (E) Embryo number of old females

(i) Effect of absolute body length across environments (zero-inflation model)

| **Fixed effect** | Estimate | *SE* | $\chi^2$ | *df* | *P* |
|---|---|---|---|---|---|
| Intercept (freshwater) | -0.804 | 0.258 | 9.685 | 1 | **0.002** |
| Environment (stable) | -0.671 | 0.378 | 3.367 | 2 | 0.186 |
| Environment (fluctuating) | -0.436 | 0.361 | | | |
| Absolute body size (standardized) | -0.738 | 0.166 | 19.883 | 1 | **<0.001** |

| **Random effect** | Variance | *sd* | Number of groups |
|---|---|---|---|
| Brood ID | 0.464 | 0.681 | 95 |

(ii) Effect of absolute body length across environments (conditional model)

| **Fixed effect** | Estimate | *SE* | $\chi^2$ | *df* | *P* |
|---|---|---|---|---|---|
| Intercept (freshwater) | 2.659 | 0.055 | 2312.072 | 1 | **<0.001** |
| Environment (stable) | 0.060 | 0.070 | 1.614 | 2 | 0.446 |
| Environment (fluctuating) | -0.025 | 0.074 | | | |
| Absolute body size (standardized) | 0.239 | 0.037 | 40.780 | 1 | **<0.001** |

| **Random effect** | Variance | *sd* | Number of groups |
|---|---|---|---|
| Brood ID | 0.034 | 0.185 | 95 |

(iii) Effect of body length within each environment (zero-inflation model)

| **Fixed effect** | Estimate | *SE* | $\chi^2$ | *df* | *P* |
|---|---|---|---|---|---|
| Intercept (freshwater) | -0.721 | 0.256 | 7.973 | 1 | **0.005** |
| Environment (stable) | -0.950 | 0.381 | 6.235 | 2 | **0.044** |
| Environment (fluctuating) | -0.376 | 0.359 | | | |
| Within-group body size (standardized) | -0.720 | 0.162 | 19.675 | 1 | **<0.001** |

| **Random effect** | Variance | *sd* | Number of groups |
|---|---|---|---|
| Brood ID | 0.453 | 0.673 | 95 |

| **Pairwise comparison** | *P* |
|---|---|
| Freshwater-Stable | **0.037** |
| Freshwater-Fluctuating | 0.580 |
| Stable-Fluctuating | 0.290 |

(iv) Effect of body length within each environment (conditional model)

| **Fixed effect** | Estimate | *SE* | $\chi^2$ | *df* | *P* |
|---|---|---|---|---|---|
| Intercept (freshwater) | 2.629 | 0.056 | 2197.316 | 1 | **<0.001** |
| Environment (stable) | 0.152 | 0.069 | 8.561 | 2 | **0.014** |
| Environment (fluctuating) | -0.030 | 0.074 | | | |
| Within-group body size (standardized) | 0.232 | 0.037 | 39.571 | 1 | **<0.001** |

| **Random effect** | Variance | *sd* | Number of groups |
|---|---|---|---|
| Brood ID | 0.034 | 0.185 | 95 |

| **Pairwise comparison** | *P* |
|---|---|
| Freshwater-Stable | 0.072 |
| Freshwater-Fluctuating | 0.914 |
| Stable-Fluctuating | **0.022** |

### (F) Likelihood of giving birth

(i) Effect of absolute body length across environments

| **Fixed effect** | Estimate | *SE* | $\chi^2$ | *df* | *P* |
|---|---|---|---|---|---|
| Intercept (freshwater) | -1.877 | 0.333 | 31.847 | 1 | **<0.001** |
| Environment (stable) | -0.124 | 0.437 | 0.740 | 2 | 0.691 |
| Environment (fluctuating) | -0.404 | 0.474 | | | |
| Absolute body size (standardized) | 0.023 | 0.195 | 0.014 | 1 | 0.906 |

| **Random effect** | Variance | *sd* | Number of groups |
|---|---|---|---|
| Brood ID | 0.317 | 0.563 | 95 |

(ii) Effect of body length within each environment

| **Fixed effect** | Estimate | *SE* | $\chi^2$ | *df* | *P* |
|---|---|---|---|---|---|
| Intercept (freshwater) | -1.878 | 0.331 | 32.294 | 1 | **<0.001** |
| Environment (stable) | -0.115 | 0.432 | 0.751 | 2 | 0.687 |
| Environment (fluctuating) | -0.404 | 0.473 | | | |
| Within-group body size (standardized) | 0.041 | 0.191 | 0.046 | 1 | 0.830 |

| **Random effect** | Variance | *sd* | Number of groups |
|---|---|---|---|
| Brood ID | 0.310 | 0.557 | 95 |

### (G) Total offspring number

(i) Effect of absolute body length across environments (zero-inflation model)

| **Fixed effect** | Estimate | *SE* | $\chi^2$ | *df* | *P* |
|---|---|---|---|---|---|
| Intercept (freshwater) | -1.012 | 0.255 | 15.687 | 1 | **<0.001** |
| Environment (stable) | -0.467 | 0.365 | 1.683 | 2 | 0.431 |
| Environment (fluctuating) | -0.260 | 0.351 | | | |
| Absolute body size (standardized) | -0.669 | 0.157 | 18.181 | 1 | **<0.001** |

| **Random effect** | Variance | *sd* | Number of groups |
|---|---|---|---|
| Brood ID | 0.135 | 0.367 | 95 |

(ii) Effect of absolute body length across environments (conditional model)

| **Fixed effect** | Estimate | *SE* | $\chi^2$ | *df* | *P* |
|---|---|---|---|---|---|
| Intercept (freshwater) | 2.651 | 0.061 | 1913.435 | 1 | **<0.001** |
| Environment (stable) | 0.120 | 0.079 | 2.568 | 2 | 0.277 |
| Environment (fluctuating) | 0.029 | 0.084 | | | |
| Absolute body size (standardized) | 0.241 | 0.040 | 36.573 | 1 | **<0.001** |

| **Random effect** | Variance | *sd* | Number of groups |
|---|---|---|---|
| Brood ID | 0.029 | 0.170 | 95 |

(iii) Effect of body length within each environment (zero-inflation model)

| **Fixed effect** | Estimate | *SE* | $\chi^2$ | *df* | *P* |
|---|---|---|---|---|---|
| Intercept (freshwater) | -0.939 | 0.252 | 13.902 | 1 | **<0.001** |
| Environment (stable) | -0.720 | 0.365 | 3.985 | 2 | 0.136 |
| Environment (fluctuating) | -0.202 | 0.350 | | | |
| Within-group body size (standardized) | -0.658 | 0.154 | 18.203 | 1 | **<0.001** |

| **Random effect** | Variance | *sd* | Number of groups |
|---|---|---|---|
| Brood ID | 0.125 | 0.354 | 95 |

(iv) Effect of body length within each environment (conditional model)

| **Fixed effect** | Estimate | *SE* | $\chi^2$ | *df* | *P* |
|---|---|---|---|---|---|
| Intercept (freshwater) | 2.621 | 0.061 | 1832.178 | 1 | **<0.001** |
| Environment (stable) | 0.213 | 0.078 | 9.418 | 2 | **0.009** |
| Environment (fluctuating) | 0.022 | 0.084 | | | |
| Within-group body size (standardized) | 0.236 | 0.039 | 36.257 | 1 | **<0.001** |

| **Random effect** | Variance | *sd* | Number of groups |
|---|---|---|---|
| Brood ID | 0.029 | 0.169 | 95 |

| **Pairwise comparison** | *P* |
|---|---|
| Freshwater-Stable | **0.018** |
| Freshwater-Fluctuating | 0.962 |
| Stable-Fluctuating | **0.043** |

**Table S7. Statistical outputs for the effects of environment, age and *body size* on male life-history and reproductive traits.** Two additional analyses were run to test if body size had an effect on the measured traits: (1) final model with absolute body length (globally standardized across all environments) as a covariate and (2) final model with within-group body length (standardized within each environment/age combination) as a covariate. Table S5 reported the the final model for each trait. *Methods* in the main text describe the details of the analysis (see *Analysis 2*).

#### (A) Relative telomere length

(i) Effect of absolute body length across environments and ages

| **Fixed effect** | Estimate | *SE* | *F* | *df* | *df.res* | *P* |
|---|---|---|---|---|---|---|
| Intercept (freshwater, old) | 0.742 | 0.043 | 294.760 | 1 | 49.714 | **<0.001** |
| Environment (stable) | -0.130 | 0.033 | 7.945 | 2 | 128.348 | **<0.001** |
| Environment (fluctuating) | -0.067 | 0.038 | | | | |
| Adult age (young) | 0.051 | 0.040 | 1.548 | 1 | 30.023 | 0.223 |
| Absolute body size (standardized) | 0.006 | 0.019 | 0.097 | 1 | 164.966 | 0.755 |

| **Random effect** | Variance | *sd* | correlation | Number of groups |
|---|---|---|---|---|
| Brood ID (intercept) | 0.033 | 0.183 | | 43 |
| Brood ID (young) | 0.019 | 0.137 | -0.870 | |
| Residual | 0.032 | 0.178 | | |

| **Pairwise comparison** | *P* |
|---|---|
| Freshwater-Stable | **<0.001** |
| Freshwater-Fluctuating | 0.194 |
| Stable-Fluctuating | 0.268 |

(ii) Effect of body length within each environment/age combination

| **Fixed effect** | Estimate | *SE* | *F* | *df* | *df.res* | *P* |
|---|---|---|---|---|---|---|
| Intercept (freshwater, old) | 0.745 | 0.042 | 314.239 | 1 | 46.920 | **<0.001** |
| Environment (stable) | -0.129 | 0.033 | 7.842 | 2 | 117.075 | **<0.001** |
| Environment (fluctuating) | -0.074 | 0.033 | | | | |
| Adult age (young) | 0.049 | 0.039 | 1.438 | 1 | 29.872 | 0.240 |
| Within-group body size (standardized) | -0.002 | 0.016 | 0.012 | 1 | 166.836 | 0.914 |

| **Random effect** | Variance | *sd* | correlation | Number of groups |
|---|---|---|---|---|
| Brood ID (intercept) | 0.032 | 0.179 | | 43 |
| Brood ID (young) | 0.019 | 0.137 | -0.870 | |
| Residual | 0.032 | 0.179 | | |

| **Pairwise comparison** | *P* |
|---|---|
| Freshwater-Stable | **<0.001** |
| Freshwater-Fluctuating | 0.066 |
| Stable-Fluctuating | 0.213 |

### (B) Immune response

(i) Effect of absolute body length across environments and ages

| **Fixed effect** | Estimate | *SE* | $\chi^2$ | *df* | *P* |
|---|---|---|---|---|---|
| Intercept (freshwater, old) | -1.902 | 0.237 | 64.426 | 1 | **<0.001** |
| Environment (stable) | 0.160 | 0.272 | 0.868 | 2 | 0.648 |
| Environment (fluctuating) | -0.144 | 0.334 | | | |
| Age (young) | -0.128 | 0.238 | 0.287 | 1 | 0.592 |
| Absolute body size (standardized) | 0.229 | 0.137 | 2.789 | 1 | 0.095 |

| **Random effect** | Variance | *sd* | correlation | Number of groups |
|---|---|---|---|---|
| Brood ID (intercept) | <0.001 | <0.001 | | 113 |
| Brood ID (young) | <0.001 | <0.001 | -0.950 | |

(ii) Effect of body length within each environment/age combination

| **Fixed effect** | Estimate | *SE* | $\chi^2$ | *df* | *P* |
|---|---|---|---|---|---|
| Intercept (freshwater, old) | -1.792 | 0.222 | 65.461 | 1 | **<0.001** |
| Environment (stable) | 0.204 | 0.271 | 4.245 | 2 | 0.120 |
| Environment (fluctuating) | -0.385 | 0.301 | | | |
| Age (young) | -0.206 | 0.234 | 0.771 | 1 | 0.380 |
| Within-group body size (standardized) | 0.198 | 0.116 | 2.923 | 1 | 0.087 |

| **Random effect** | Variance | *sd* | correlation | Number of groups |
|---|---|---|---|---|
| Brood ID (intercept) | <0.001 | <0.001 | | 113 |
| Brood ID (young) | <0.001 | <0.001 | -0.900 | |

### (C) Number of mating attempts

Due to a significant environment*age interaction, body size effects were were tested for separately at each age:

(i) Young males: effect of absolute body length across environments (zero-inflation model)

| **Fixed effect** | Estimate | *SE* | $\chi^2$ | *df* | *P* |
|---|---|---|---|---|---|
| Intercept (freshwater) | -3.235 | 0.633 | 26.077 | 1 | **<0.001** |
| Environment (stable) | 0.067 | 0.811 | 1.515 | 2 | 0.469 |
| Environment (fluctuating) | -1.702 | 1.493 | | | |
| Absolute body size (standardized) | -0.043 | 0.429 | 0.010 | 1 | 0.921 |

| **Random effect** | Variance | *sd* | Number of groups |
|---|---|---|---|
| Brood ID (intercept) | <0.001 | 0.001 | 98 |

(ii) Young males: effect of absolute body length across environments (conditional model)

| **Fixed effect** | Estimate | *SE* | $\chi^2$ | *df* | *P* |
|---|---|---|---|---|---|
| Intercept (freshwater) | 3.843 | 0.090 | 1829.300 | 1 | **<0.001** |
| Environment (stable) | -0.095 | 0.113 | 0.911 | 2 | 0.634 |
| Environment (fluctuating) | -0.099 | 0.123 | | | |
| Absolute body size (standardized) | -0.078 | 0.053 | 2.142 | 1 | 0.143 |

| **Random effect** | Variance | *sd* | Number of groups |
|---|---|---|---|
| Brood ID (intercept) | 0.015 | 0.123 | 98 |

(iii) Young males: effect of body length within each environment (zero-inflation model)

| **Fixed effect** | Estimate | *SE* | $\chi^2$ | *df* | *P* |
|---|---|---|---|---|---|
| Intercept (freshwater) | -3.254 | 0.608 | 28.615 | 1 | **<0.001** |
| Environment (stable) | 0.067 | 0.810 | 1.617 | 2 | 0.446 |
| Environment (fluctuating) | -1.658 | 1.413 | | | |
| Within-group body size (standardized) | -0.075 | 0.381 | 0.039 | 1 | 0.844 |

| **Random effect** | Variance | *sd* | Number of groups |
|---|---|---|---|
| Brood ID (intercept) | <0.001 | 0.001 | 98 |

(iv) Young males: effect of body length within each environment (conditional model)

| **Fixed effect** | Estimate | *SE* | $\chi^2$ | *df* | *P* |
|---|---|---|---|---|---|
| Intercept (freshwater) | 3.809 | 0.088 | 1894.134 | 1 | **<0.001** |
| Environment (stable) | -0.095 | 0.113 | 0.911 | 2 | 0.634 |
| Environment (fluctuating) | -0.012 | 0.109 | | | |
| Within-group body size (standardized) | -0.071 | 0.045 | 2.499 | 1 | 0.114 |

| **Random effect** | Variance | *sd* | Number of groups |
|---|---|---|---|
| Brood ID (intercept) | 0.016 | 0.125 | 98 |

(v) Old males: effect of absolute body length across environments (zero-inflation model)

| **Fixed effect** | Estimate | *SE* | $\chi^2$ | *df* | *P* |
|---|---|---|---|---|---|
| Intercept (freshwater) | -18.736 | 2540.147 | <0.001 | 1 | 0.994 |
| Environment (stable) | 15.777 | 2540.147 | 1.462 | 2 | 0.482 |
| Environment (fluctuating) | 14.616 | 2540.147 | | | |
| Absolute body size (standardized) | -0.605 | 0.471 | 1.652 | 1 | 0.199 |

(vi) Old males: effect of absolute body length across environments (conditional model)

| **Fixed effect** | Estimate | *SE* | $\chi^2$ | *df* | *P* |
|---|---|---|---|---|---|
| Intercept (freshwater) | 3.104 | 0.087 | 1276.808 | 1 | **<0.001** |
| Environment (stable) | 0.322 | 0.118 | 12.145 | 2 | **0.002** |
| Environment (fluctuating) | 0.377 | 0.121 | | | |
| Absolute body size (standardized) | 0.037 | 0.055 | 0.460 | 1 | 0.498 |

| **Random effect** | Variance | *sd* | Number of groups |
|---|---|---|---|
| Brood ID (intercept) | 0.015 | 0.122 | 94 |

| **Pairwise comparison** | *P* |
|---|---|
| Freshwater-Stable | **0.019** |
| Freshwater-Fluctuating | **0.006** |
| Stable-Fluctuating | 0.905 |

(vii) Old males: effect of body length within each environment (zero-inflation model)

| **Fixed effect** | Estimate | *SE* | $\chi^2$ | *df* | *P* |
|---|---|---|---|---|---|
| Intercept (freshwater) | -19.905 | 3361.946 | <0.001 | 1 | 0.995 |
| Environment (stable) | 16.628 | 3361.946 | 0.323 | 2 | 0.851 |
| Environment (fluctuating) | 16.133 | 3361.946 | | | |
| Within-group body size (standardized) | -0.606 | 0.427 | 2.018 | 1 | 0.155 |

(viii) Old males: effect of absolute body length within each environment (conditional model)

| **Fixed effect** | Estimate | *SE* | $\chi^2$ | *df* | *P* |
|---|---|---|---|---|---|
| Intercept (freshwater) | 3.115 | 0.086 | 1325.238 | 1 | **<0.001** |
| Environment (stable) | 0.330 | 0.117 | 11.345 | 2 | **0.003** |
| Environment (fluctuating) | 0.339 | 0.112 | | | |
| Within-group body size (standardized) | 0.037 | 0.046 | 0.653 | 1 | 0.419 |

| **Random effect** | Variance | *sd* | Number of groups |
|---|---|---|---|
| Brood ID (intercept) | 0.015 | 0.121 | 94 |

| **Pairwise comparison** | *P* |
|---|---|
| Freshwater-Stable | **0.014** |
| Freshwater-Fluctuating | **0.008** |
| Stable-Fluctuating | 0.997 |

#### (D) Likelihood of successful mating

Due to a significant environment*age interaction, body size effects were were tested for separately at each age:

(i) Effect of absolute body length across environments (young males)

| Fixed effect | Estimate | *SE* | $\chi^2$ | *df* | *P* |
|---|---|---|---|---|---|
| Intercept (freshwater) | -1.172 | 0.275 | 18.210 | 1 | **<0.001** |
| Environment (stable) | -0.022 | 0.356 | 0.019 | 2 | 0.991 |
| Environment (fluctuating) | 0.030 | 0.395 | | | |
| Absolute body size (standardized) | 0.087 | 0.167 | 0.273 | 1 | 0.601 |

| Random effect | Variance | *sd* | Number of groups |
|---|---|---|---|
| Brood ID (intercept) | <0.001 | <0.001 | 98 |

(ii) Effect of body length within each environment (young males)

| Fixed effect | Estimate | *SE* | $\chi^2$ | *df* | *P* |
|---|---|---|---|---|---|
| Intercept (freshwater) | -1.135 | 0.264 | 18.476 | 1 | **<0.001** |
| Environment (stable) | -0.022 | 0.356 | 0.040 | 2 | 0.980 |
| Environment (fluctuating) | -0.067 | 0.349 | | | |
| Within-group body size (standardized) | 0.082 | 0.141 | 0.340 | 1 | 0.560 |

| Random effect | Variance | *sd* | Number of groups |
|---|---|---|---|
| Brood ID (intercept) | <0.001 | <0.001 | 98 |

#### (iii) Effect of absolute body length across environments (old males)

| **Fixed effect** | Estimate | *SE* | $\chi^2$ | *df* | *P* |
|---|---|---|---|---|---|
| Intercept (freshwater) | -1.983 | 0.448 | 19.633 | 1 | **<0.001** |
| Environment (stable) | -0.941 | 0.552 | 3.017 | 2 | 0.221 |
| Environment (fluctuating) | -0.443 | 0.527 | | | |
| Absolute body size (standardized) | -0.068 | 0.256 | 0.071 | 1 | 0.789 |

| **Random effect** | Variance | *sd* | Number of groups |
|---|---|---|---|
| Brood ID (intercept) | 1.073 | 1.036 | 94 |

#### (iv) Effect of body length within each environment (old males)

| **Fixed effect** | Estimate | *SE* | $\chi^2$ | *df* | *P* |
|---|---|---|---|---|---|
| Intercept (freshwater) | -2.000 | 0.438 | 20.833 | 1 | **<0.001** |
| Environment (stable) | -0.956 | 0.548 | 3.052 | 2 | 0.217 |
| Environment (fluctuating) | -0.377 | 0.461 | | | |
| Within-group body size (standardized) | -0.075 | 0.212 | 0.124 | 1 | 0.725 |

| **Random effect** | Variance | *sd* | Number of groups |
|---|---|---|---|
| Brood ID (intercept) | 1.063 | 1.031 | 94 |

### (E) Time spent with female

(i) Effect of absolute body length across environments and ages

| **Fixed effect** | Estimate | *SE* | *F* | *df* | *df.res* | *P* |
|---|---|---|---|---|---|---|
| Intercept (freshwater, old) | 364.209 | 17.815 | 413.265 | 1 | 229.152 | **<0.001** |
| Environment (stable) | 12.515 | 20.333 | 0.662 | 2 | 303.513 | 0.516 |
| Environment (fluctuating) | 25.483 | 22.031 | | | | |
| Adult age (young) | -49.013 | 15.886 | 9.426 | 1 | 89.481 | **0.003** |
| Absolute body size (standardized) | 9.703 | 9.624 | 0.990 | 1 | 283.385 | 0.321 |

| **Random effect** | Variance | *sd* | correlation | Number of groups |
|---|---|---|---|---|
| Brood ID (intercept) | 4697 | 68.540 | | 98 |
| Brood ID (young) | 6970 | 83.490 | -0.970 | |
| Male ID (intercept) | 7195 | 84.830 | | 323 |
| Residual | 21670 | 147.210 | | |

(ii) Effect of body length within each environment/age combination

| **Fixed effect** | Estimate | *SE* | *F* | *df* | *df.res* | *P* |
|---|---|---|---|---|---|---|
| Intercept (freshwater, old) | 368.903 | 17.190 | 455.679 | 1 | 218.685 | **<0.001** |
| Environment (stable) | 13.570 | 20.311 | 0.344 | 2 | 287.994 | 0.710 |
| Environment (fluctuating) | 15.424 | 19.642 | | | | |
| Adult age (young) | -51.506 | 15.698 | 10.657 | 1 | 84.346 | **0.002** |
| Within-group body size (standardized) | 7.196 | 8.126 | 0.765 | 1 | 291.295 | 0.383 |

| **Random effect** | Variance | *sd* | correlation | Number of groups |
|---|---|---|---|---|
| Brood ID (intercept) | 4684 | 68.440 | | 98 |
| Brood ID (young) | 6960 | 83.430 | -0.970 | |
| Male ID (intercept) | 7196 | 84.830 | | 323 |
| Residual | 21675 | 147.220 | | |

**(F) Sperm velocity- VCL**

Due to a significant environment*age interaction, body size effects were tested for separately at each age:

(i) Effect of absolute body length across environments (young males)

| **Fixed effect** | Estimate | *SE* | *F* | *df* | *df.res* | *P* |
|---|---|---|---|---|---|---|
| Intercept (freshwater) | 176.797 | 2.837 | 3865.889 | 1 | 253.380 | **<0.001** |
| Environment (stable) | -7.396 | 3.403 | 3.031 | 2 | 240.790 | 0.0501 |
| Environment (fluctuating) | -7.980 | 3.815 | | | | |
| Absolute body size (standardized) | 4.445 | 1.753 | 6.335 | 1 | 264.190 | **0.012** |

| **Random effect** | Variance | *sd* | Number of groups |
|---|---|---|---|
| Brood ID (intercept) | 128.000 | 11.310 | 97 |
| Residual | 444.800 | 21.090 | |

(ii) Effect of body length within each environment (young males)

| **Fixed effect** | Estimate | *SE* | *F* | *df* | *df.res* | *P* |
|---|---|---|---|---|---|---|
| Intercept (freshwater) | 178.671 | 2.749 | 4209.156 | 1 | 251.320 | **<0.001** |
| Environment (stable) | -7.336 | 3.409 | 7.445 | 2 | 224.320 | **<0.001** |
| Environment (fluctuating) | -12.928 | 3.342 | | | | |
| Within-group body size (standardized) | 3.448 | 1.474 | 5.394 | 1 | 264.940 | **0.021** |

| **Random effect** | Variance | *sd* | Number of groups |
|---|---|---|---|
| Brood ID (intercept) | 127.700 | 11.300 | 97 |
| Residual | 446.700 | 21.140 | |

| **Pairwise comparison** | *P* |
|---|---|
| Freshwater-Stable | 0.083 |
| Freshwater-Fluctuating | **<0.001** |
| Stable-Fluctuating | 0.168 |

(iii) Effect of absolute body length across environments (old males)

| **Fixed effect** | Estimate | *SE* | *F* | *df* | *df.res* | *P* |
|---|---|---|---|---|---|---|
| Intercept (freshwater) | 166.243 | 2.357 | 4939.958 | 1 | 225.190 | **<0.001** |
| Environment (stable) | 1.087 | 3.140 | 0.511 | 2 | 259.020 | 0.601 |
| Environment (fluctuating) | 3.373 | 3.337 | | | | |
| Absolute body size (standardized) | 2.642 | 1.523 | 2.969 | 1 | 270.070 | 0.086 |

| **Random effect** | Variance | *sd* | Number of groups |
|---|---|---|---|
| Brood ID (intercept) | 69.000 | 8.307 | 94 |
| Residual | 380.300 | 19.501 | |

(iv) Effect of body length within each environment (old males)

| **Fixed effect** | Estimate | *SE* | *F* | *df* | *df.res* | *P* |
|---|---|---|---|---|---|---|
| Intercept (freshwater) | 167.015 | 2.315 | 5169.176 | 1 | 225.060 | **<0.001** |
| Environment (stable) | 1.696 | 3.124 | 0.147 | 2 | 245.480 | 0.864 |
| Environment (fluctuating) | 0.788 | 2.989 | | | | |
| Within-group body size (standardized) | 1.928 | 1.290 | 2.205 | 1 | 271.110 | 0.139 |

| **Random effect** | Variance | *sd* | Number of groups |
|---|---|---|---|
| Brood ID (intercept) | 69.740 | 8.351 | 94 |
| Residual | 380.960 | 19.518 | |

#### (G) Total sperm count (log-transformed)

(i) Effect of absolute body length across environments and ages

| **Fixed effect** | Estimate | *SE* | *F* | *df* | *df.res* | *P* |
|---|---|---|---|---|---|---|
| Intercept (freshwater, old) | 15.342 | 0.059 | 66170.329 | 1 | 191.074 | **<0.001** |
| Environment (stable) | -0.163 | 0.059 | 5.851 | 2 | 277.129 | **0.003** |
| Environment (fluctuating) | -0.201 | 0.065 | | | | |
| Adult age (young) | -0.309 | 0.065 | 22.692 | 1 | 93.456 | **<0.001** |
| Absolute body size (standardized) | 0.175 | 0.029 | 35.487 | 1 | 294.719 | **<0.001** |

| **Random effect** | Variance | *sd* | correlation | Number of groups |
|---|---|---|---|---|
| Brood ID (intercept) | 0.115 | 0.340 | | 97 |
| Brood ID (young) | 0.217 | 0.466 | -0.930 | |
| Male ID (intercept) | 0.029 | 0.170 | | 312 |
| Residual | 0.206 | 0.454 | | |

| **Pairwise comparison** | *P* |
|---|---|
| Freshwater-Stable | **0.017** |
| Freshwater-Fluctuating | **0.006** |
| Stable-Fluctuating | 0.833 |

(ii) Effect of body length within each environment/age combination

| **Fixed effect** | Estimate | *SE* | *F* | *df* | *df.res* | *P* |
|---|---|---|---|---|---|---|
| Intercept (freshwater, old) | 15.433 | 0.058 | 71267.891 | 1 | 176.308 | **<0.001** |
| Environment (stable) | -0.152 | 0.059 | 24.388 | 2 | 260.903 | **<0.001** |
| Environment (fluctuating) | -0.389 | 0.056 | | | | |
| Adult age (young) | -0.359 | 0.065 | 30.135 | 1 | 90.338 | **<0.001** |
| Within-group body size (standardized) | 0.147 | 0.024 | 36.593 | 1 | 301.590 | **<0.001** |

| **Random effect** | Variance | *sd* | correlation | Number of groups |
|---|---|---|---|---|
| Brood ID (intercept) | 0.119 | 0.345 | | 97 |
| Brood ID (young) | 0.232 | 0.481 | -0.940 | |
| Male ID (intercept) | 0.030 | 0.173 | | 312 |
| Residual | 0.203 | 0.451 | | |

| **Pairwise comparison** | *P* |
|---|---|
| Freshwater-Stable | **0.029** |
| Freshwater-Fluctuating | **<0.001** |
| Stable-Fluctuating | **<0.001** |

**Table S8. Statistical outputs for effects of developmental environment and adult age on shape deformity of (A) females and (B) males**

**(A) Females**

(i) Initial model with environment*age interactive effect

| **Fixed effect** | Estimate | *SE* | $\chi^2$ | *df* | *P* |
|---|---|---|---|---|---|
| Intercept (freshwater, old) | -14.193 | 2.611 | 29.550 | 1 | **<0.001** |
| Environment (stable) | -0.143 | 3.364 | 0.115 | 2 | 0.944 |
| Environment (fluctuating) | 0.758 | 2.907 | | | |
| Age (young) | -15.621 | 4.358 | 12.851 | 1 | **<0.001** |
| Environment (stable) * Age (young) | 0.254 | 5.375 | 0.003 | 2 | 0.999 |
| Environment (fluctuating) * Age (young) | 0.034 | 4.745 | | | |

| **Random effect** | Variance | *sd* | Number of groups |
|---|---|---|---|
| Brood ID (intercept) | 0.029 | 0.171 | 95 |
| Female ID (intercept) | 6468.000 | 80.426 | 284 |

(ii) Final model excluding environment*age interaction to interpret the main effects

| **Fixed effect** | Estimate | *SE* | $\chi^2$ | *df* | *P* |
|---|---|---|---|---|---|
| Intercept (freshwater, old) | -14.196 | 2.611 | 29.563 | 1 | **<0.001** |
| Environment (stable) | -0.136 | 3.357 | 0.115 | 2 | 0.944 |
| Environment (fluctuating) | 0.759 | 2.906 | | | |
| Age (young) | -15.544 | 2.911 | 28.514 | 1 | **<0.001** |

| **Random effect** | Variance | *sd* | Number of groups |
|---|---|---|---|
| Brood ID (intercept) | 0.001 | 0.029 | 95 |
| Female ID (intercept) | 6472.000 | 80.447 | 284 |

We found no effect of the developmental environment on female skeletal deformity, but old females ($n$ = 32 of 284) were more often deformed than young females ($n$ = 15 of 284).

### (B) Males

(i) Initial model with environment*age interactive effect

| **Fixed effect** | Estimate | *SE* | $\chi^2$ | *df* | *P* |
|---|---|---|---|---|---|
| Intercept (freshwater, old) | -14.280 | 2.760 | 26.778 | 1 | **<0.001** |
| Environment (stable) | 0.063 | 3.435 | 0.004 | 2 | 0.998 |
| Environment (fluctuating) | 0.184 | 3.221 | | | |
| Age (young) | -14.748 | 3.972 | 13.786 | 1 | **<0.001** |
| Environment (stable) * Age (young) | -1.966 | 6.265 | 0.326 | 2 | 0.850 |
| Environment (fluctuating) * Age (young) | 1.389 | 4.383 | | | |

| **Random effect** | Variance | *sd* | Number of groups |
|---|---|---|---|
| Brood ID (intercept) | 36.610 | 6.050 | 94 |
| Male ID (intercept) | 6924.650 | 83.210 | 277 |

(ii) Final model excluding environment*age interaction to interpret the main effects

| **Fixed effect** | Estimate | *SE* | $\chi^2$ | *df* | *P* |
|---|---|---|---|---|---|
| Intercept (freshwater, old) | -14.382 | 2.768 | 26.995 | 1 | **<0.001** |
| Environment (stable) | -0.021 | 3.523 | 0.007 | 2 | 0.997 |
| Environment (fluctuating) | 0.210 | 3.227 | | | |
| Age (young) | -15.114 | 2.889 | 27.361 | 1 | **<0.001** |

| **Random effect** | Variance | *sd* | Number of groups |
|---|---|---|---|
| Brood ID (intercept) | 41.790 | 6.464 | 94 |
| Male ID (intercept) | 7550.570 | 86.894 | 277 |

We found no effect of the developmental environment on male skeletal deformity, but old males ($n$ = 30 of 277) were more often deformed than young males ($n$ = 19 of 277).

**Table S9. Statistical outputs (Procrustes multivariate analysis of variances) of developmental environment and age effects on body shape**

(A) Model with the environment*age interaction for *female* shape

| | *df* | SS | MS | *F* | *P* |
|---|---|---|---|---|---|
| Log centroid size | 1 | 0.043 | 0.043 | 96.174 | **0.001** |
| Environment | 2 | 0.006 | 0.003 | 6.369 | **0.001** |
| Age | 1 | 0.008 | 0.008 | 18.203 | **0.001** |
| Environment/Brood | 174 | 0.127 | <0.001 | 7.963 | **0.001** |
| Environment*Age | 2 | <0.001 | <0.001 | 0.012 | 0.491 |
| Environment/Brood*Age | 174 | 0.066 | <0.001 | -2.056 | 0.981 |
| Residuals | 149 | 0.066 | <0.001 | | |
| total | 503 | 0.316 | | | |

(B) Model with the individual main effects of environment and age for *female* shape

| | *df* | SS | MS | *F* | *P* |
|---|---|---|---|---|---|
| Log centroid size | 1 | 0.043 | 0.043 | 104.194 | **0.001** |
| Environment | 2 | 0.006 | 0.003 | 6.900 | **0.001** |
| Age | 1 | 0.008 | 0.008 | 19.720 | **0.001** |
| Environment/Brood | 174 | 0.127 | <0.001 | 1.778 | **0.001** |
| Residuals | 325 | 0.133 | <0.001 | | |
| total | 503 | 0.316 | | | |

The developmental environment and adult age independently affected a female shape. Females from the fluctuating salinity environment had a more ventral mouth and upper insertion of the anal fin than females from the other two environments (Supplementary Fig. 9). Old females had more dorsally positioned eyes, a smaller head/body ratio and a deeper abdomen than young females (Supplementary Fig. 9).

(C) Models with the environment*age interaction for *male* shape

| | *df* | SS | MS | *F* | *P* |
|---|---|---|---|---|---|
| Log centroid size | 1 | 0.017 | 0.016 | 31.340 | **0.001** |
| Environment | 2 | 0.010 | 0.005 | 9.116 | **0.001** |
| Age | 1 | 0.032 | 0.032 | 61.567 | **0.001** |
| Environment /brood | 183 | 0.165 | 0.001 | 1.714 | **0.001** |
| Environment*Age | 2 | 0.003 | 0.002 | 3.173 | **0.001** |
| Environment/Brood*Age | 183 | 0.085 | <0.001 | 0.887 | 0.967 |
| Residuals | 121 | 0.064 | <0.001 | | |
| total | 493 | 0.376 | | | |

For males, the effect of the developmental environment on body shape changed with age (i.e., an interaction). Young males from the fluctuating salinity environment had a narrower body with a more ventral mouth than males from the other two environments, but these shape differences were reduced in old males (Supplementary Fig. 10).

**Table S10. Pairwise Procrustes distances between least squares means for shapes of (A) females (with the significant environment effect) and (B) males (with the significant interaction between environment and adult age).**

(A) Pairwise comparisons of the environment effect for *female* shape

| | F | SS | FS |
|---|---|---|---|
| Freshwater (F) | | 0.001 | 0.001 |
| Stable salinity (SS) | 0.006 | | 0.003 |
| Fluctuating salinity (FS) | 0.006 | 0.005 | |

(B) Pairwise comparisons of the environment and age interaction for *male* shape

| | Old F | Young F | Old SS | Young SS | Old FS | Young FS |
|---|---|---|---|---|---|---|
| Old F | | 0.001 | 0.002 | --- | 0.001 | --- |
| Young F | 0.018 | | --- | 0.002 | --- | 0.001 |
| Old SS | 0.009 | --- | | 0.001 | 0.003 | --- |
| Young SS | --- | 0.008 | 0.019 | | --- | 0.035 |
| Old FS | 0.010 | --- | 0.008 | --- | | 0.001 |
| Young FS | --- | 0.013 | --- | 0.007 | 0.014 | |

Note: Absolute differences of distances between least squares mean are below the diagonal, and their associated *P* values are above the diagonal (based on randomized residual permutation procedure with 1000 permutations).

**Table S11. Statistical outputs for the effects of developmental environment, adult age and their interaction on size-corrected gonopodium length.**

(A) Model with environment*age interactive effect

| **Fixed effect** | Estimate | *SE* | *F* | *df* | *df.res* | *P* |
|---|---|---|---|---|---|---|
| Intercept (freshwater, old) | 1.860 | 0.004 | 211043 | 1 | 279.69 | **<0.001** |
| Environment (stable) | -0.014 | 0.005 | 28.075 | 2 | 790.32 | **<0.001** |
| Environment (fluctuating) | -0.037 | 0.005 | | | | |
| Age (young) | 0.004 | 0.004 | 1.381 | 1 | 249.61 | 0.241 |
| Log-transformed absolute body size (standardized) | 0.049 | 0.002 | 871.033 | 1 | 638.47 | **<0.001** |
| Environment (stable) * Age (young) | 0.007 | 0.005 | 10.297 | 2 | 339.08 | **<0.001** |
| Environment (fluctuating) * Age (young) | 0.020 | 0.004 | | | | |

| **Random effect** | Variance | *sd* | correlation | Number of groups |
|---|---|---|---|---|
| Brood ID (intercept) | <0.001 | 0.021 | | 116 |
| Brood ID (young) | <0.001 | 0.014 | -0.69 | |
| Male ID (intercept) | 0.001 | 0.023 | | 628 |
| Residual | <0.001 | 0.022 | | |

Due to a significant environment*age interaction, the environmental effect was tested for seperately at each age:

(B) young males

| **Fixed effect** | Estimate | *SE* | *F* | *df* | *df.res* | *P* |
|---|---|---|---|---|---|---|
| Intercept (freshwater) | 1.860 | 0.003 | 413010 | 1 | 313.357 | **<0.001** |
| Environment (stable) | -0.007 | 0.003 | 8.781 | 2 | 612.97 | **<0.001** |
| Environment (fluctuating) | -0.016 | 0.004 | | | | |
| Log-transformed absolute body size (standardized) | 0.052 | 0.002 | 911.870 | 1 | 607.01 | **<0.001** |

| **Random effect** | Variance | *sd* | Number of groups |
|---|---|---|---|
| Brood ID (intercept) | <0.001 | 0.016 | 116 |
| Residual | <0.001 | 0.030 | |

| **Pairwise comparison** | *P* |
|---|---|
| Freshwater-Stable | 0.082 |
| Freshwater-Fluctuating | **<0.001** |
| Stable-Fluctuating | **0.049** |

(C) Old males

| **Fixed effect** | Estimate | *SE* | *F* | *df* | *df.res* | *P* |
|---|---|---|---|---|---|---|
| Intercept (freshwater) | 1.869 | 0.004 | 171624.601 | 1 | 223.37 | **<0.001** |
| Environment (stable) | -0.020 | 0.006 | 14.214 | 2 | 256.50 | **<0.001** |
| Environment (fluctuating) | -0.033 | 0.006 | | | | |
| Log-transformed absolute body size (standardized) | 0.051 | 0.003 | 316.238 | 1 | 272.03 | **<0.001** |

| **Random effect** | Variance | *sd* | Number of groups |
|---|---|---|---|
| Brood ID (intercept) | <0.001 | 0.017 | 94 |
| Residual | 0.001 | 0.036 | |

| **Pairwise comparison** | *P* |
|---|---|
| Freshwater-Stable | **0.003** |
| Freshwater-Fluctuating | **<0.001** |
| Stable-Fluctuating | 0.100 |

The length of male intromittent organ (i.e., gonopodium) increased with size (Supplementary Fig. 11). Correcting for body size, the developmental environment and adult age interacted to affect gonopodium length. Size-corrected gonopodium length was significantly shorter in young males from the fluctuating salinity environment than those from the freshwater ($P <$ 0.001) or stable salinity environments ($P = 0.049$), who did not differ from each other ($P =$ 0.082). When old, however, males from both salinity environments ($P = 0.100$) had shorter size-corrected gonopodium than freshwater males (both $P < 0.004$).
